# DHHC3 mediated Cadm4 palmitoylation regulates myelination in CNS

**DOI:** 10.1101/2022.09.23.509146

**Authors:** Yanli Chang, Jiangli Zhu, Chenchen Nie, Yajuan Lu, Fangjing Ren, Xize Cao, Juanjuan Li, Changhong Wang, Chenyu Yang, Tianhan Li, Yinming Liang, Shiqian Qi, Xiaohong Kang, Eryan Kong

**Affiliations:** The Second Affiliated Hospital of Xinxiang Medical University, Xinxiang, China; Institute of Psychiatry and Neuroscience, Xinxiang key laboratory of protein palmitoylation and major human diseases, Xinxiang Medical University, Xinxiang, China; Department of Urology, State Key Laboratory of Biotherapy and Cancer Center, West China Hospital, Sichuan University and National Collaborative Innovation Center, Chengdu, 610041, China; Center of Cryo-Electron Microscopy, Zhejiang University, Hangzhou, China; Laboratory of Genetic Regulators in the Immune System, Henan Key Laboratory of Immunology and Targeted Therapy, School of Laboratory Medicine, Xinxiang Medical University, Xinxiang, China; Department of Oncology, the First Affiliated Hospital of Xinxiang Medical University, Xinxiang, China

**Author notes:** equally contributde (Yanli Chang, Jiangli Zhu). Correspondence: Dr. Eryan Kong, or.

**Keywords:** Cadm4, protein palmitoylation, DHHC3, membrane stability, myelination

## Abstract

Cell adhesion molecule 4 (Cadm4) plays important roles on plasma membrane (PM) to regulate myelin formation and the downregulation of Cadm4 is a prominent feature in many demyelination diseases. However, how Cadm4 maintains its level on PM has been elusive. Here, we identify that Cadm4 is palmitoylated at cysteine-347 (C347) and palmitoylation regulates the stable localization of Cadm4 on PM, as blocking palmitoylation by mutating C347 into alanine (C347A) results in the dissociation of Cadm4 from PM and targeting for degradation. Intriguingly, blocking Cadm4 palmitoylation by introducing C347A (Cadm4-KI) causes myelin abnormalities in CNS, characterized by loss of myelination, myelin infoldings and hypermyelination. Moreover, it is uncovered that Cadm4 palmitoylation is catalyzed by DHHC3, reducing Cadm4 palmitoylation by the deletion of DHHC3 renders the redistribution of Cadm4 for degrading. Consistently, the genetic deletion of DHHC3 leads to downregulated Cadm4 palmitoylation and defects in CNS myelination, virtually phenocopies that of the Cadm4-KI mice. Our findings suggest a mechanism that the stable localization of Cadm4 on PM regulated by protein palmitoylation is vital for myelination in CNS.

## Introduction

Myelination is a fundamental feature of the nervous system, which attributes the living animals the ability to take swift actions in response to emergency (1, 2). The saltatory propagation of the action potentials transmitted within the nerve cells is guaranteed by the myelin sheath(3), a specific organization of axonal/glia membranes and numerous molecules along the axonal shaft including ion-channel proteins and cell adhesion molecules (Cadms)(3, 4). The formation of the myelin sheath involves the cellular events that the membrane of glia (oligodendrocyte in CNS or Schwann cell in PNS) spirals around the axonal membrane, which requires the reciprocal interactions of cell surface proteins, in particular, the Cadms(5–7). Cadms have 4 members in its family, of which Cadm 1-3 can express both on glia and axon, while Cadm4 is mainly expressed in glia cells (5, 7). The heterophilic bindings of Cadm2/3 and Cadm4 were shown to be the most common interactions observed(7, 8).

Cadms, also known as synaptic cell adhesion molecules (SynCAM) or Nectin-like adhesion molecules (Necl), are featured by a single trans-membrane domain, an extracellular Ig-like domains and a short cytoplasmic tail(1, 2). The heterophilic interactions between Cadms are mediated by the Ig-like domains, the intracellular tail of Cadms contains motifs that can bind protein 4.1G and PDZ sequence(9, 10). While it seems logic to assume that the single trans-membrane domain of Cadms is sufficient for its stable localization on the plasma membrane (PM), this assumption has not been examined. Moreover, it was suggested that the proper level of Cadms on PM should be precisely maintained as either increased or decreased expression of Cadm4/Cadm3 inhibits proper myelination (5, 6). Interestingly, it was revealed that decreased level of Cadm4 is a striking feature in many demyelination diseases including Charcot-Marie-Tooth neuropathy(11). Yet, how the stable localization and the level of Cadm4 regulated remains largely unexplored.

Of note, to identify palmitoylated proteins in the central nervous system (CNS) by palm-proteomics in our lab(12), we spotted Cadm4, implying that Cadm4 is palmitoylated (Fig. S1A). Protein S-palmitoylation can be reversibly modified by adding (catalyzed by DHHC1-24) or removing palmitate (Catalyzed by APT1/2, PPT1/2 and ABHD17a) on/off from cysteine residue(12–16), which substantially alter the hydrophobicity of the modified protein and thus regulate its membrane localization, protein-protein interaction and downstream signaling(15, 17, 18). Interestingly, understanding why Cadm4 should be palmitoylated *in vivo* lead us to uncover that DHHC3 mediated palmitoylation stabilizes Cadm4 and therefore regulates its level on PM, which is vital for proper myelination in CNS.

## Results

### Cadm4 is palmitoylated at cysteine-347

To verify if Cadm4 is modified with protein palmitoylation, Cadm4 expressed either ectopically in N2a, a neuroblastoma cell line, or endogenously in WT mice brain was evaluated by Acyl-RAC assay(12, 19). The results showed that Cadm4 is readily palmitoylated both *in vitro* and *in vivo* (Fig. 1A–1B). For confirmation, 2-BP (a general inhibitor of protein palmitoylation) was used to incubate with cells expressing Cadm4-Flag, the experiments showed that 2-BP could effectively reduce the level of palmitoylated-Cadm4 (palm-Cadm4) in a time and dose dependent manner (Fig. 1C–1D). Curiously, we noticed that multiple bands of Cadm4 appear in WB when Cadm4-Flag is expressed either in N2a or HEK-293T cells (Fig. 1A, 1C–1D and Fig. S1B), interestingly, PNG treatment (an inhibitor of protein glycosylation) could flatten these bands into a single band (Fig. S1C).

**Fig. 1.**
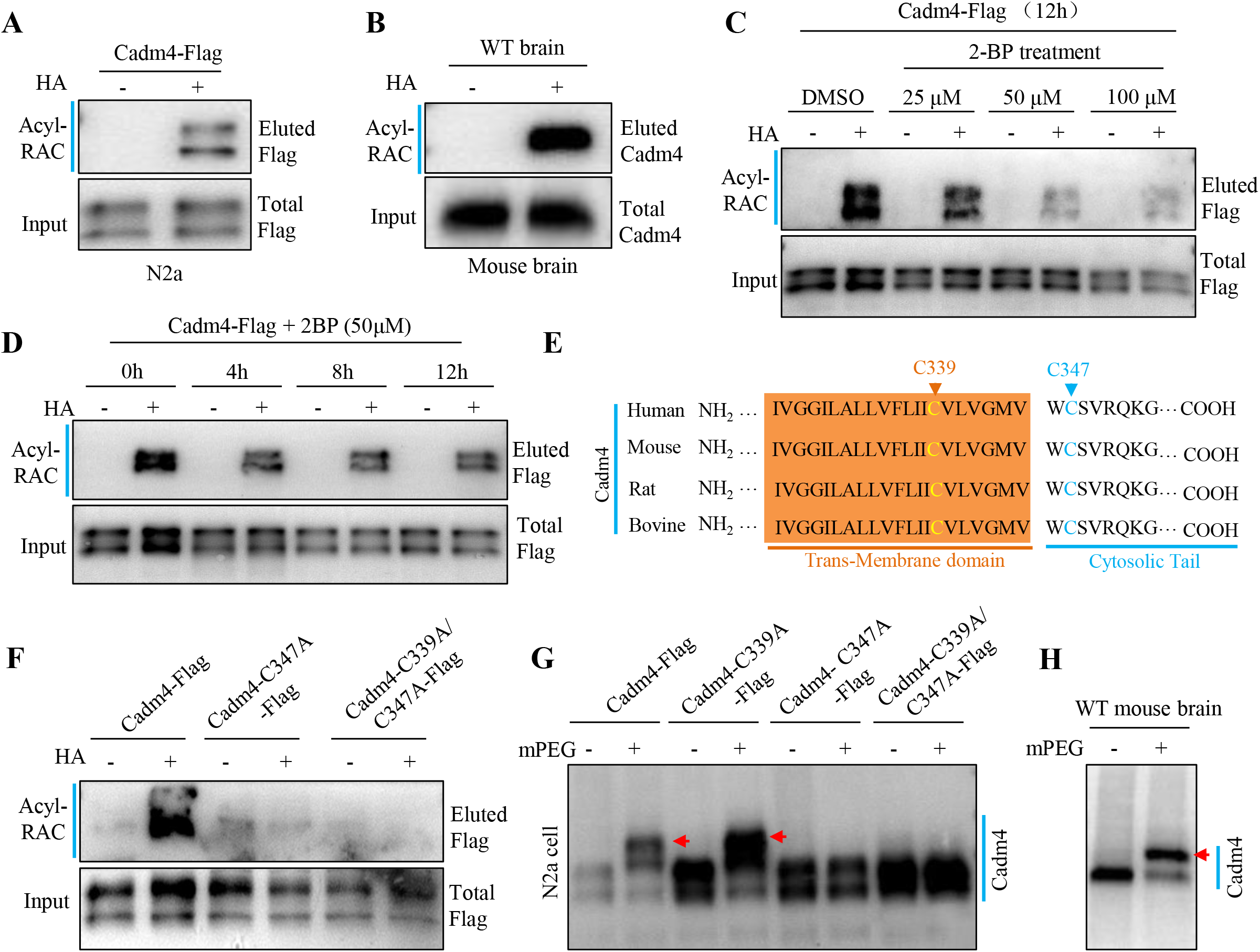
Cadm4 is palmitoylated at cysteine-347. **A-B**, Cadm4 expressed in N2a cells or from mice brain was analyzed for protein palmitoylation by Acyl-RAC assay. HA+, with hydroxylamine, HA-, without hydroxylamine. **C-D**, N2a cells expressing Cadm4-Flag was incubated with 2-BP for various dose and time, and were evaluated for the level of palm-Cadm4. **E**, Protein sequences of Cadm4 from various species were aligned for analyzing cysteine conservation. **F**, Cadm4 or its mutants (C347A and C339A/C347A) were expressed in N2a cells and subjected for Acyl-RAC assay. **G-H**, Cadm4 and its mutants (C347A and C339A/C347A) expressed in N2a cells (**G**) or endogenously expressed Cadm4 (**H**) were processed with mPEG-labeling assay. The mPEG labeling causes the band shift, pointed with red arrows.

To identify the cysteine residue that is palmitoylated, we analyzed the protein sequence of Cadm4 by intentionally looking at its transmembrane (TM) domain, considering that palmitoylated proteins often associate with hydrophobic membranes(15, 18). The analysis showed that cysteine-339 (C339) and C347 are close to the boundary of the TM domain and conserved in different species (Fig. 1E). Accordingly, point mutations were introduced into Cadm4. The Acyl-RAC assay showed that while Cadm4 is palmitoylated, the single mutation in C347 (C347A) is sufficient to block its palmitoylation (Fig. 1F and Fig. S1D), suggesting that C347 is the major site of palmitoylation in Cadm4. To confirm this finding, the mPEG labeling assay, where palmitate is substituted by mPEG(14), was performed to show that the C347A mutation could fully block the labeling of mPEG-Cadm4 (displayed by no migrated band), yet the WT Cadm4 or Cadm4-C339A can be labeled by mPEG (single migrated band, Fig. 1G and S1E). The single migrated mPEG-Cadm4 band (arrow pointed, Fig. 1G) suggests that Cadm4 is likely modified with single site of palmitoylation. Furthermore, the mPEG assay demonstrated that endogenously expressed Cadm4 is also palmitoylated at single site since only one migrated mPEG-Cadm4 band was visualized (arrow pointed, Fig. 1H). Together, these data showed that Cadm4 is palmitoylated at C347.

### Palmitoylation regulates Cadm4 for PM localization

To test if palmitoylation is involved in regulating Cadm4 for PM localization, Cadm4-GFP and Cadm4-C347A-GFP were expressed in N2a cells. It showed that as Cadm4 resides mainly on PM, a large proportion of Cadm4-C347A distributes within the cytosol as vesicle-like structures (Fig. 2A–2B), suggesting that blocking palmitoylation somehow weakens the PM localization of Cadm4. Similarly, the incubation of 2-BP, aiming at reducing the level of palm-Cadm4, with N2a cells expressing Cadm4-GFP redistributes Cadm4 from PM to cytosol (Fig. 2C–2D). Noticeably, it may or may not be relevant that as Cadm4 (with Cadm4-C347A expressing or upon 2-BP treatment) dissociates from PM into cytosol, often, the PM marker protein Na/K-ATPase loses its membrane localization and appears in cytosol as puncta shape (a sign of internalization) as well. Moreover, we prepared the plasma membrane fractions (Fig. S1F) of cultured N2a cells, the results showed that reducing the level of palm-Cadm4 either by C347A mutation or 2-BP significantly lowers the level of Cadm4 in the membrane fraction as compare to that of the control (WT-Cadm4) (Fig. 2E–2F). Hereby, these data suggest that palmitoylation is involved in regulating Cadm4 for PM localization.

**Fig. 2.**
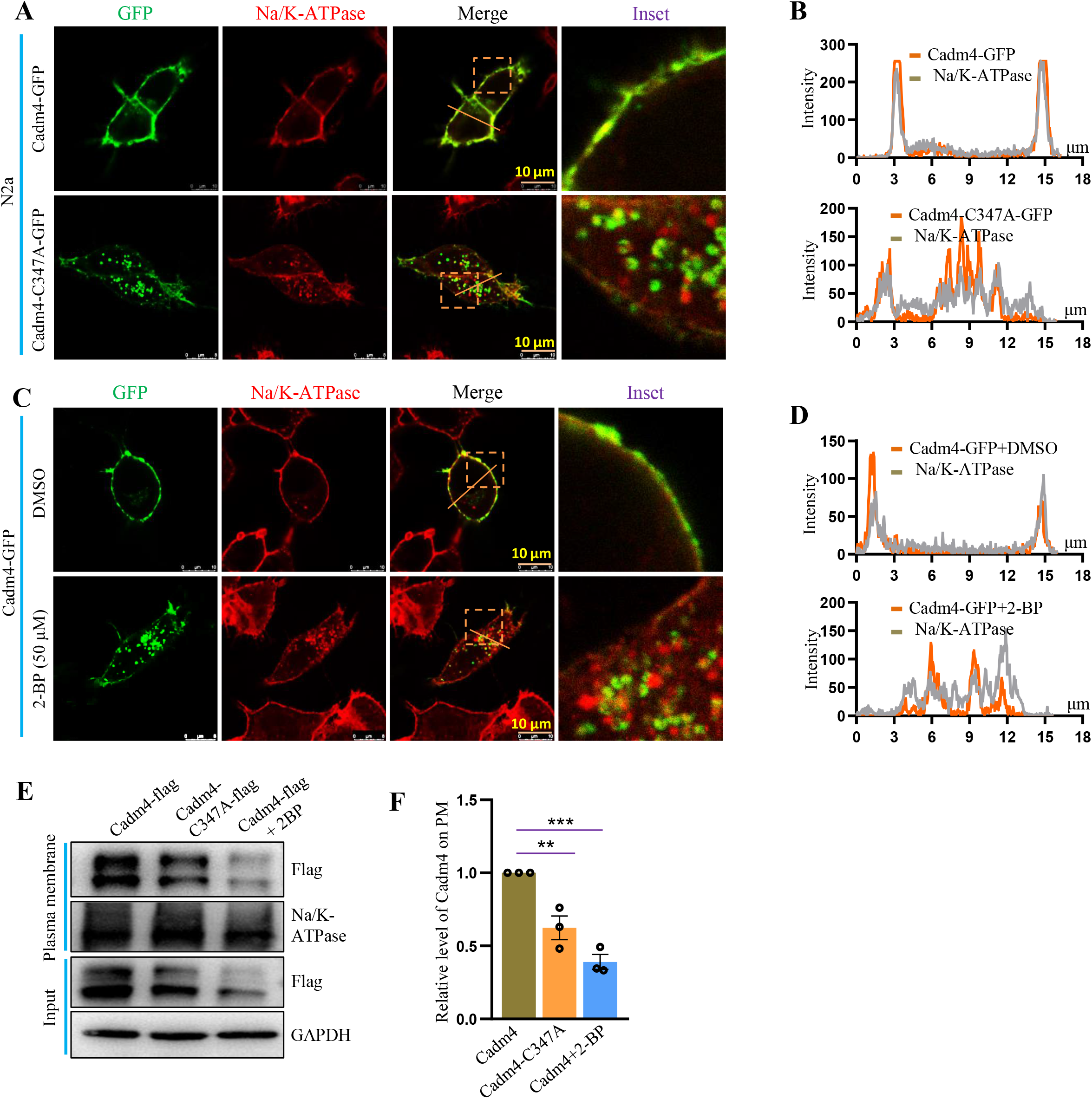
Palmitoylation regulates Cadm4 for PM localization. **A**, Cadm4 and Cadm4-C347A were expressed in N2a cells and fixed for immunofluorescence analysis, Na/K-ATPase is a marker for plasma membrane. **B**, The profile of the intensity of the red and green fluorescence was measured along the slash. **C**, N2a cells expressing Cadm4 were incubated with DMSO or 2-BP (50μM), and fixed for immunofluorescence analysis. **D**, The profile of the intensity of the red and green fluorescence was measured along the slash. **E-F**, N2a cells expressing Cadm4 or Cadm4-C347A were treated with DMSO or 2-BP for the preparation of plasma membrane fractions, which were evaluated (**E**) and quantified (**F**). *P*=0.001, n=3 biological replicates, one-way ANOVA; Bonferroni post hoc test comparing Cadm4 and Cadm4-C347A, \*\**P*=0.009; Cadm4 and Cadm4+2-BP, \*\*\**P*= 0.001. Data are mean ± s.e.m.

### Depalmitoylated Cadm4 is prone to be internalized from PM for degradation

To understand the nature of the vesicle-like structures encircling Cadm4 when palm-Cadm4 is downregulated by either C347A mutation or 2-BP treatment (Fig. 2A and 2C), RFP-Lamp1 (lysosome marker) or -Rab7 (late endosome) was expressed for colocalization. The analysis showed that the colocalization rate of Cadm4 with Lamp1 or Rab7 is significantly upregulated when Cadm4-C347A is expressed or upon the treatment of 2-BP (Fig. 3A–3B and S2A–S2B), implying that depalmitoylated Cadm4 might be internalized into vesicles and scheduled for degradation. To test this assumption, we thought to examine if palmitoylation regulates the stability of Cadm4 as well, cycloheximide (CHX) was used to block *de novo* protein synthesis in N2a cells. Interestingly, the experiments showed that reducing the level of palm-Cadm4 (C347A mutation or 2-BP treatment) significantly enhances the degradation of Cadm4 as compared to that of the control (Fig. 3C–3D and 3E–3F). To identify the potential pathway that is involved in degrading Cadm4, inhibitors as CHQ (Chloroquine, blocking lysosome degradation) and 3MA (3-Methyladenine, blocking autophagy) were applied. It showed that either 3-MA or CHQ could markedly inhibit the degradation of Cadm4-C347A (rectangle circled lane, Fig. 3G), indicating that an autophagy-lysosome pathway is likely engaged for degrading Cadm4-C347A.

**Fig. 3.**
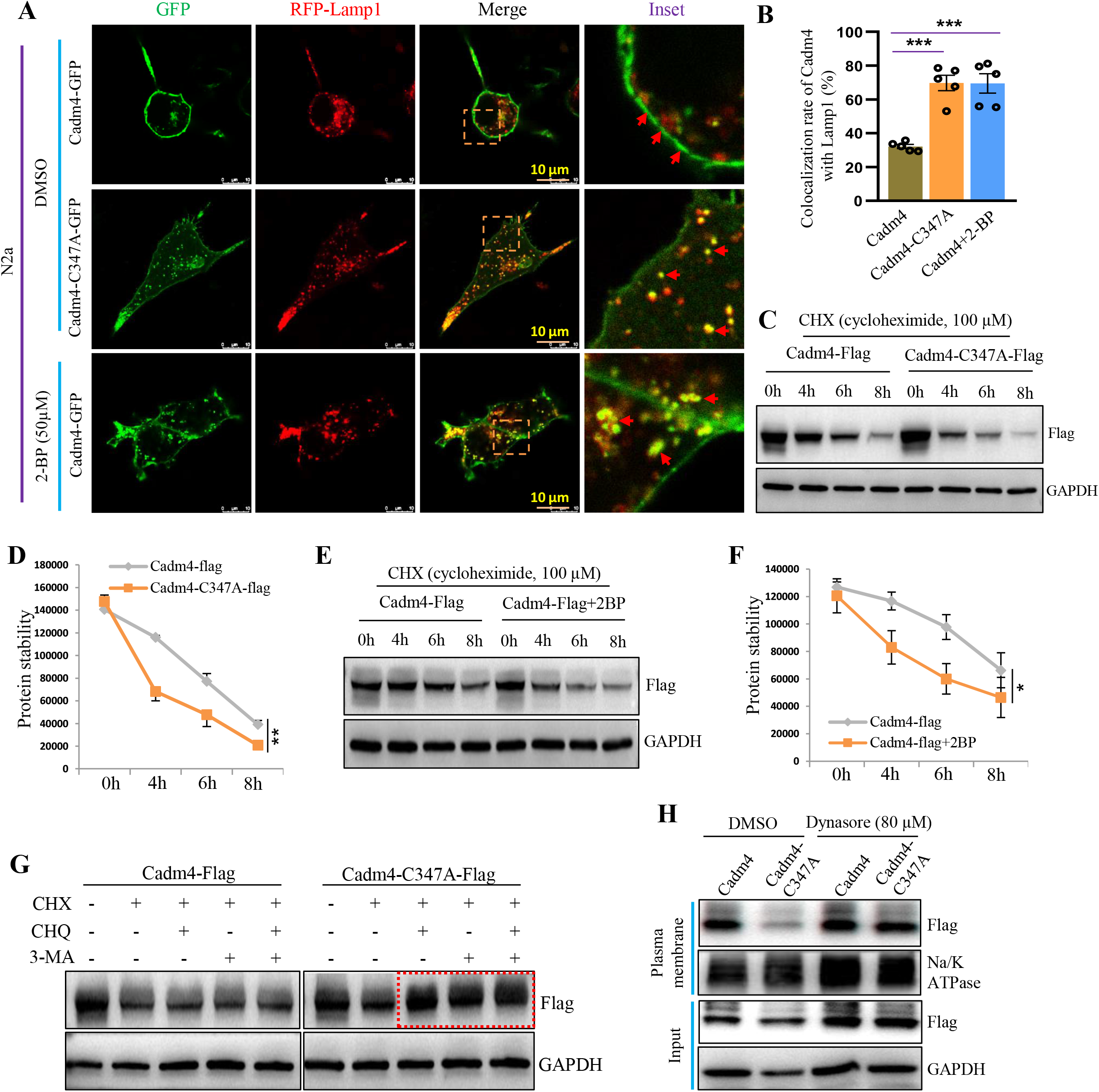
Depalmitoylated Cadm4 is prone to be internalized from PM for degradation. **A**, Cadm4 or Cadm4-C347A was expressed in N2a cells, treated with DMSO or 2-BP (50 μM) and fixed for immunofluorescence analysis, RFP-Lamp1 was used to label lysosome. **B**, The colocalization rate of Cadm4 and Lamp1 was quantified. *P*< 0.001, n=5 cells, one-way ANOVA; Bonferroni post hoc test comparing Cadm4 and Cadm4-C347A, \*\*\**P*<0.001; Cadm4 and Cadm4+2-BP, \*\*\**P*<0.001. **C-D**, N2a cells expressing Cadm4 or Cadm4-C347A were incubated with CHX (cycloheximide, 100μM) for various length of time, subjected for WB analysis (**C**) and the levels of Cadm4 were quantified (**D**). \*\**P*=0.007, t-test (2-tailed, n=3 biological replicates). **E-F**, N2a cells expressing Cadm4 treated with DMSO or 2-BP (50 μM) were incubated with CHX for different periods, analyzed by WB (**E**) and the levels of Cadm4 were quantified (**F**). \*\**P*=0.018, t-test (2-tailed, n=3 biological replicates). **G**, N2a cells expressing Cadm4 or Cadm4-C347A were incubated with or without CHX, CHQ (Chloroquine, 50 μM) or 3-MA (Methyladenine, 3mM), and subjected for WB analysis. **H**, The plasma membrane fractions were prepared from N2a cells expressing Cadm4 or Cadm4-C347A, treated with either DMSO or Dynasore (80 μM). Data are mean ± s.e.m.

To briefly explore how Cadm4-C347A is internalized from the PM, Dynasore (mainly blocking the clathrin-mediated endocytosis) was incubated with N2a cells expressing Cadm4-C347A. The results showed that the treatment of Dynasore could recover the level of Cadm4-C347A on the PM as that of the WT Cadm4 (Fig. 3H and Fig. S2C), indicating that palmitoylation is not essential for its PM localization but is important for the maintenance of its stable PM localization. Above all, these evidences implied that as Cadm4 is depalmitoylated, it is prone to be internalized from PM and scheduled for degradation via the autophagy and lysosome path.

### Blocking Cadm4 palmitoylation *in vivo* leads to severe defects in myelination

As depalmitoylated Cadm4 is prone to be internalized and the PM presence of Cadm4 is crucial for myelination, one might speculate that blocking Cadm4 palmitoylation *in vivo* might interfere myelin formation. To test this idea, a point mutation (C347A) was introduced into Cadm4 to block Cadm4 palmitoylation in C57BL/B6 mice (hereafter named Cadm4-KI) (Fig. S3A–3C). The Cadm4-KI mice seems normal at birth in general.

Firstly, we verified that Cadm4 is indeed expressed principally in oligodendrocytes rather than neurons (Fig. S4A), and Cadm4 palmitoylation is inhibited in Cadm4-KI as compared to that of the WT mice (Fig. 4A). Then, 3-months old mice (notably, the expression of Cadm4 also peaks at 3M, Fig. S4B) were sacrificed for analyzing the status of myelination in corpus callosum by electron microscopy (Fig. 4B). The results showed that the myelinated axons per field of view is dramatically downregulated (Fig. 4I), while the percentage of abnormal myelination (loss of myelination (Fig. 4C), myelin infoldings (Fig. 4D), vacuolation (Fig. 4E), detachment of axon from myelin sheath (Fig. 4F), thickened myelin sheath/hypermyelination (Fig. 4G) and myelination contains multiple axons (Fig. 4H)) is significantly upregulated in Cadm4-KI as compared to that of the WT mice (Fig. 4J). Furthermore, the value of g-ratio is significantly reduced in Cadm4-KI as comparing to WT mice (Fig. 4K), mainly due to the fact that a proportion of the myelin sheath is markedly thickened in Cadm4-KI (Red arrows pointed, Fig. 4B) mice.

**Fig. 4.**
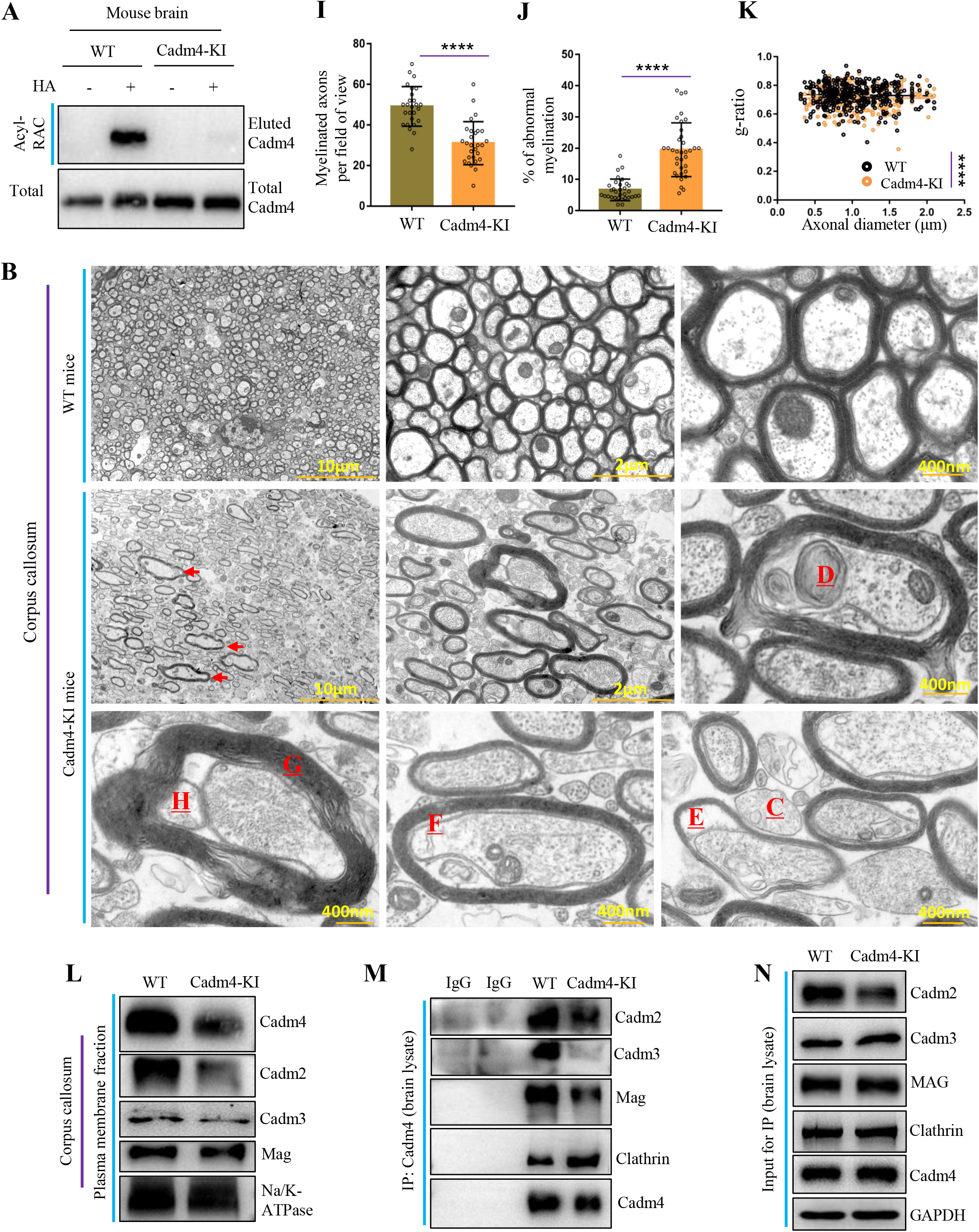
Blocking Cadm4 palmitoylation *in vivo* leads to severe defects in myelination. **A**, Brain lysates from WT and Cadm4-KI mice were evaluated for the level of palm-Cadm4. **B-H**, Corpus callosum was isolated from prefixed brains of WT and Cadm4-KI mice and examined for myelination under electron microscopy. Abnormal myelination is pointed out by red arrows or labeled: **C**, loss of myelination, **D**, myelin infoldings, **E**, vacuolation, **F**, detachment of axon from myelin sheath, **G**, thickened myelin sheath/hypermyelination, **H**, myelination contains multiple axons. **I-K**, Myelinated axons per field of view (**I**, \*\*\*\**P*<0.0001, t-test), % of abnormal myelination (**J**, \*\*\*\**P*<0.0001, t-test, 2-tailed), and g-ratio (**K**, \*\*\*\**P*<0.0001, t-test, 2-tailed) were quantified from 3-6 mice. **L**, The plasma membrane fractions were prepared from the corpus callosum of WT and Cadm4-KI mice and analyzed by WB. **M**, Immunoprecipitation was performed with the brain lysates of WT and Cadm4-KI mice by using Cadm4 antibody, the IP eluent was analyzed by WB for Cadm4 binding proteins. Data are mean ± s.e.m.

To explore if such phenotype correlates with the downregulated level of Cadm4 on PM in Cadm4-KI mice, the PM fraction of corpus collosum was prepared and the results showed that the level of Cadm4 is clearly downregulated in Cadm4-KI as compared to that of the WT mice (Fig. 4L). Moreover, as Cadm4 forms heterodimers with other cell adhesion molecules e.g. Cadm2/3 and Mag to regulate myelin formation, we also evaluated their levels on PM and the data showed that the localizations of Cadm2/3 and Mag on PM are also impaired in Cadm4-KI mice as compared to that of the WT control (Fig. 4L). Furthermore, it was shown that the heterophilic interactions between Cadm4-Cadm2/3 and -Mag are indeed undermined *in vivo*, interestingly, the binding of Cadm4 with Clathrin is increased in Cadm4-KI mice as compared to that of the WT control (Fig. 4M–4N). Together, these results tend to support the scenario that blocking Cadm4 palmitoylation reduces its localization and interactions with Cadm2/3 and Mag on PM, meanwhile may enhance its binding with Clathrin for internalization, jointly they cause severe defects in CNS myelination.

### DHHC3 palmitoylates Cadm4 and regulates its PM localization and stability

To identify the potential enzyme that palmitoylates Cadm4, all DHHCs were coexpressed with Cadm4 in N2a cells for the evaluation of palm-Cadm4, the results showed that DHHC3 could strongly enhance the level of palm-Cadm4. And, we noted that the mRNA level of DHHC3 is well expressed in N2a and oligodendrocytes (Fig. S4D–S4E). For verification, we constructed the DHHC3-KO (Using the strategy of disrupting enzyme activity center as in Fig. S5; due to the problem of lacking specificity of the DHHC3-antibody, the evidence at its protein level is not shown) cells in N2a, the Acyl-RAC assay showed that the deletion of DHHC3 diminishes the level of palm-Cadm4 (Fig. 5A). Next, we thought to examine if the membrane localization of Cadm4 is affected in DHHC3-KO cells, the images showed that as Cadm4 concentrates on PM in WT cells, a large proportion of Cadm4 relocates into cytosol as vesicle-like structures in DHHC3-KO cells (Fig. 5B–5D). Similarly, these vesicles are colocalized with lysosome marker Lamp1 (Fig. 5F–5G), as shown earlier in Fig. 3A, indicating that they are scheduled for degradation. Additionally, the membrane fractions were prepared and the results showed that upon the deletion of DHHC3, Cadm4 is apparently downregulated in PM fraction as compared to that of the control (Fig. 5E).

**Fig. 5.**
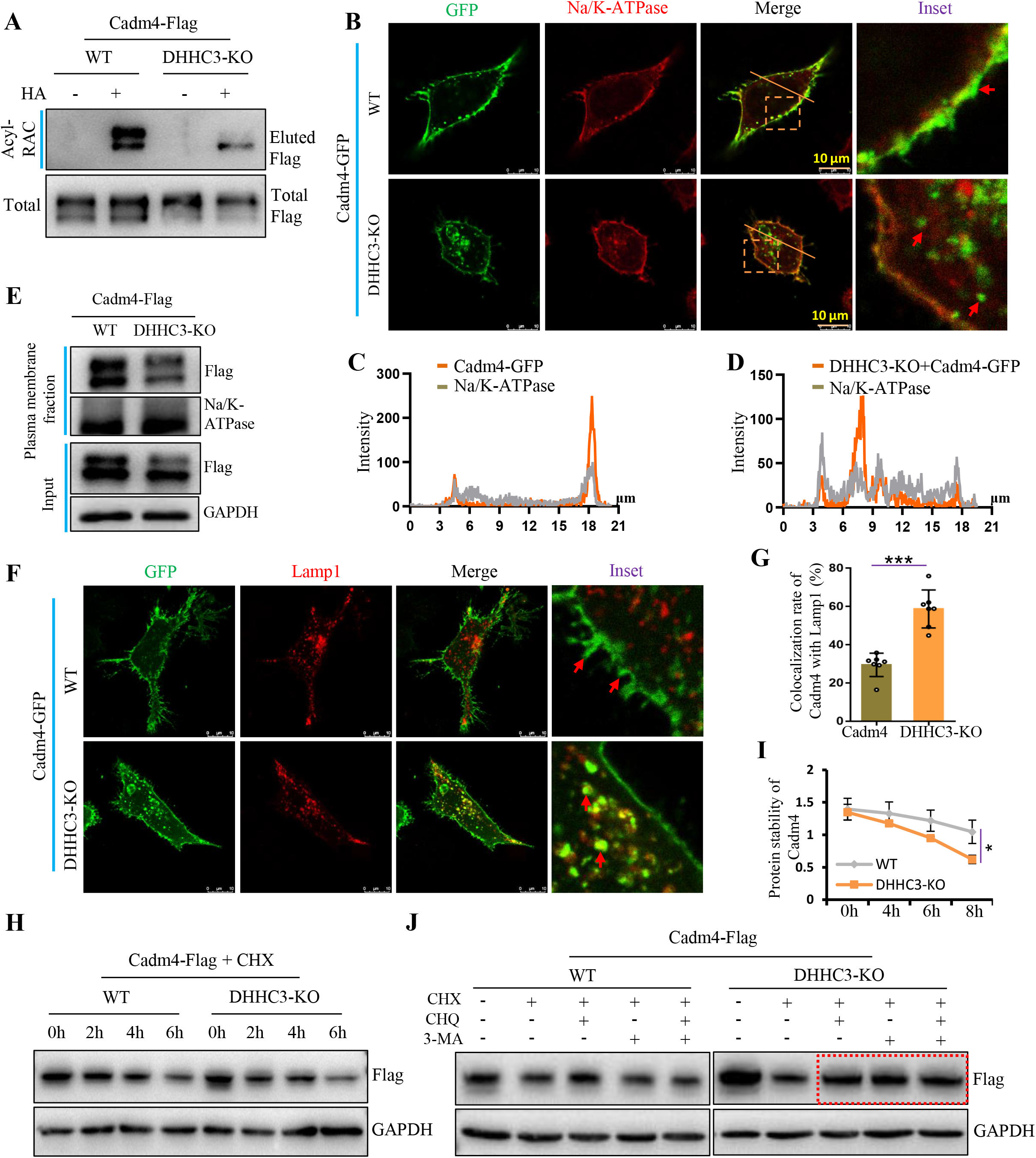
DHHC3 palmitoylates Cadm4 and regulates its PM localization and stability. **A**, Cadm4-Flag was expressed in DHHC3-KO and WT N2a cells and evaluated for protein palmitoylation by Acyl-RAC assay. **B**, Cadm4-GFP was expressed in WT and DHHC3-KO cells, and fixed for immunofluorescence analysis, Na/K-ATPase is a marker for plasma membrane. **C-D**, The profile of the intensity of the red and green fluorescence was measured along the slash. **E**, The plasma membrane fractions were prepared from WT or DHHC3-KO cells expressing Cadm4, and subjected for WB analysis. **F-G**, Cadm4-GFP was expressed in WT and DHHC3-KO cells and fixed for immunofluorescence analysis (**F**), Lamp1 is a marker for lysosome; accordingly, the colocalization rate of Cadm4 with Lamp1 was quantified (**G**). \*\*\**P*<0.001, t-test (2-tailed, n=7 cells) **H-I**, WT or DHHC3-KO cells expressing Cadm4-Flag were incubated with CHX (100μM) for different period, subjected for WB analysis (**H**) and quantified (**I**). \**P*<0.05, t-test (2-tailed, n=3 biological replicates). **J**, WT or DHHC3-KO cells expressing Cadm4-Flag were incubated with or without CHX, CHQ (50 μM) or 3-MA (3mM) and subjected for WB analysis. Data are mean ± s.e.m.

Consequently, it was conceived that the removal of DHHC3 might also affect the stability of Cadm4. The CHX-treatment experiments showed that as DHHC3 is absent, the degradation of Cadm4 is accelerated (Fig. 5H–5I). Consistently, such accelerated protein degradation can be inhibited by the treatment of 3-MA or CHQ, suggesting of the autophagy and lysosome path involved (Fig. 5J). Jointly, the above data strengthened the notion that Cadm4 palmitoylation, mediated by DHHC3, is crucial for its localization on PM and protein stability.

### DHHC3 mediated Cadm4 palmitoylation is vital for proper myelination in CNS

To further extend the evidence that Cadm4 palmitoylation is important for myelination in CNS (Fig. 5B and S4C), we reasoned that downregulating palm-Cadm4 by deleting DHHC3 *in vivo* might affect myelin formation as well. To verify, DHHC3-KO mice were generated (Fig. S5A–5C) and it showed that the removal of DHHC3 lowers the level of palm-Cadm4 in the brain of DHHC3-KO as compared to that of WT mice (Fig. 6A). Moreover, to evaluate the status of myelination in corpus collosum, images of the electron microcopy (Fig. 6B) showed that as the myelinated axons per field of view is significantly downregulated (Fig. 6H), the percentage of abnormal myelination (loss of myelination (Fig. 6C), myelin infoldings (Fig. 6D), vacuolation (Fig. 6E), axonal atrophy or detachment of axon from myelin sheath (Fig. 6F) and thickened myelin sheath (Fig. 6G)) is dramatically upregulated in DHHC3-KO as compared to that of the WT mice (Fig. 6I), almost phenocopies that of the Cadm4-KI mice (Fig. 4B). Similarly, the value of g-ratio is also downregulated in DHHC3-KO as compared to that of the WT mice (Fig. 6J), caused by the similar reason that part of the myelin sheath is thickened in DHHC3-KO mice (Red arrows pointed, Fig. 6B).

**Fig. 6.**
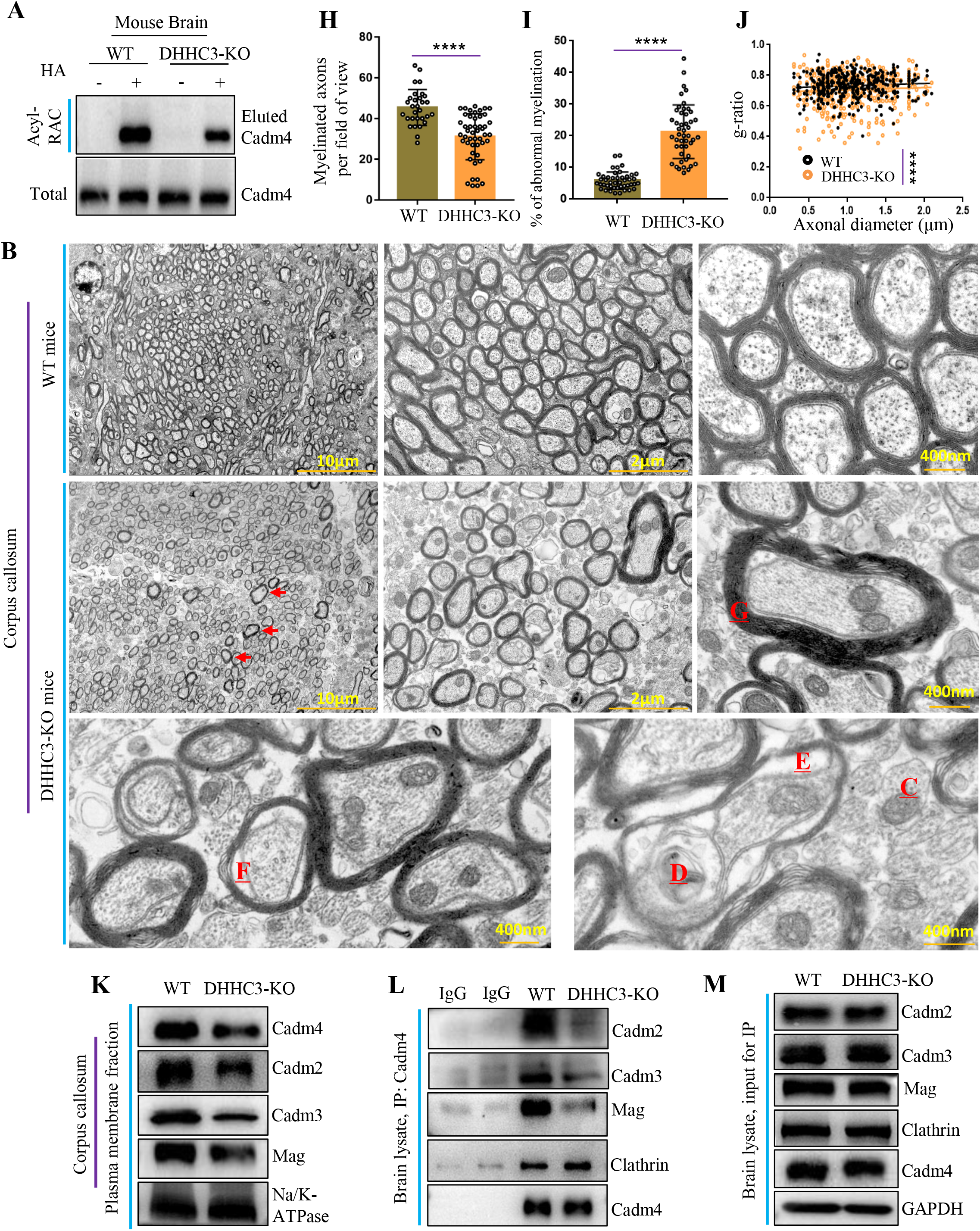
DHHC3 mediated Cadm4 palmitoylation is vital for proper myelination in CNS. **A**, Brain lysates from WT and DHHC3-KO mice were evaluated for the level of palm-Cadm4 by Acyl-RAC assay. **B-G**, Corpus callosum was isolated from prefixed brains of WT and DHHC3-KO mice and examined for myelination under electron microscopy. Abnormal myelination is pointed out by red arrows or labeled: **C**, loss of myelination, **D**, myelin infoldings, **E**, vacuolation, **F**, detachment of axon from myelin sheath, **G**, thickened myelin sheath/hypermyelination. **H-J**, Myelinated axons per field of view (**H**, \*\*\*\**p*< 0.0001, t-test), % of abnormal myelination (**I**, \*\*\*\**p*< 0.0001, t-test) and g-ratio (**J**, \*\*\*\**p*< 0.0001, t-test) were quantified from 3-6 mice. **K**, The plasma membrane fractions were prepared from the corpus callosum of WT and DHHC3-KO mice, and analyzed by WB. **L-M**, Immunoprecipitation were performed with the brain lysates of WT and DHHC3-KO mice by using Cadm4 antibody, the IP eluent was then analyzed by WB for Cadm4 binding proteins. Data are mean ± s.e.m.

Furthermore, we showed that as DHHC3 is absent, downregulated palm-Cadm4 lowers its localization on PM, as well as Cadm2/3 and Mag (Fig. 6K). Possibly, as a result, this dampens the interactions of Cadm4-Cadm2/3 and Cadm4-Mag, while augment the Cadm4-Clathrin interaction in DHHC3-KO as compared to that of the WT mice (Fig. 6L–6M). In total, these data tend to support that DHHC3 mediated palmitoylation regulates proper myelination via controlling the PM presence of Cadm4 in CNS.

## Discussion

It was shown that the axon-glia contact mediated by the surface proteins (Cadms and Mag) on PM are vital for myelination(5–8, 11). Among these, Cadm4 plays a unique role, as Cadm4 is predominantly expressed in oligodendrocytes in CNS but very scarce in neurons(5, 6) (Fig. S4A). Therefore, the strength of the axon-glia contact depends on the interactions between Cadm4 and Cadm2-3/Mag on the surfaces(20, 21), which points out that the presence and the level of Cadm4 on PM might be delicately regulated, yet the mechanism was unclear. Here, we showed that DHHC3 mediated palmitoylation maintains the stable localization of Cadm4 on PM and ensures proper myelination; reducing Cadm4 palmitoylation by either point mutation (C347A) or deleting DHHC3 somehow enhances the interaction of Cadm4-Clathrin for internalization/degradation and thus weakens the binding between Cadm4 and Cadm2-3/Mag on the surface, which results in varied myelin abnormalities (Fig. 7A).

**Fig. 7.**
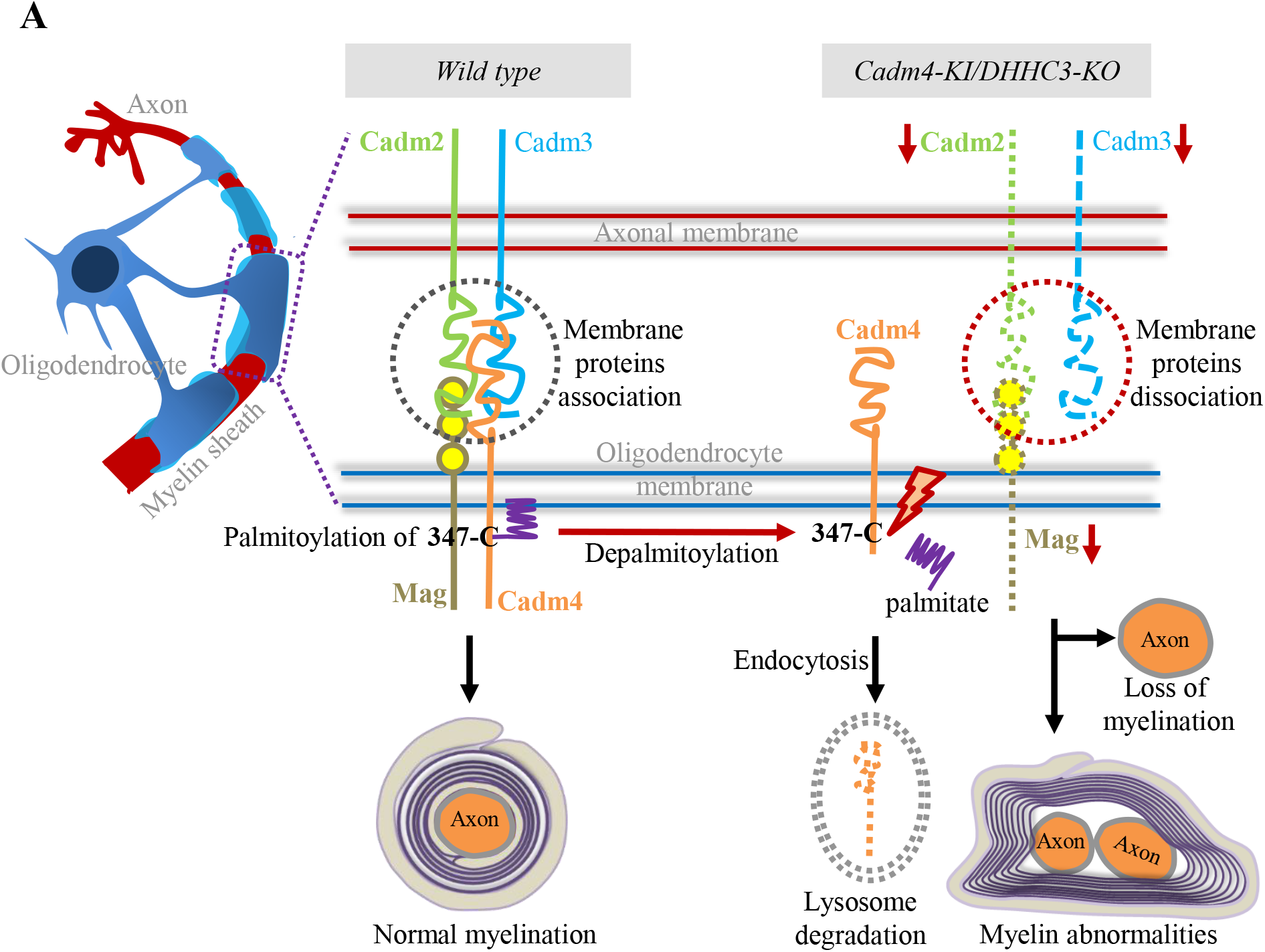
The proposed mechanism that DHHC3 mediated palmitoylation targets Cadm4 for membrane localization and regulates myelination in CNS. Cadm4 is palmitoylated at cysteine-347 to sustain its stable localization and protein level on PM, which is required for keeping its binding affinity with Cadm2/3 and Mag and ensures the proper formation of myelin sheath in CNS. As the level of palm-Cadm4 is downregulated either by Cadm4-C347A mutation or the disruption of DHHC3, the localization and the level of Cadm4 on PM is alleviated due to internalization and degradation, which impairs the surface protein interactions, weakens the binding of the axonal/glial membranes, and might contribute to myelin abnormalities in CNS.

During myelination, Oligodendrocytes send multiple exploratory cellular processes that contact numerous axons, some stabilize and develop into myelin sheath while the others retract(22, 23), indicating that myelination is a dynamic process and it is plausible that the underlying machinery regulating myelination might adapt to this dynamicity as well. Indeed, we presented here in principal that DHHC3-regulated Cadm4 palmitoylation (Fig. 5), a reversible process, is engaged in controlling the formation of myelin sheath in CNS (Fig. 6). Considering that the deletion of DHHC3 only partially downregulates the level of palm-Cadm4 (Fig. 5A and Fig. 6A), implying that other DHHCs might be also involved in palmitoylating DHHC3; moreover, to complete the cycle of palmitoylation and depalmitoylation, it will be interesting to determine which enzyme (APT1/2, PPT1/2 and ABHD17a) might catalyzes Cadm4 depalmitoylation.

By analyzing palmitoylated proteins containing single/multiple TM domain(s), it is found out that these proteins tend to be modified at the cysteine residue close to the TM or cytoplasmic boundary(24–28), interestingly, such rule is applicable to Cadm4 as Cadm4 is palmitoylated at cysteine-347, a region right after the TM domain and in the beginning of the cytoplasmic tail (Fig. 1E). Palmitoylation at such specific site might have different functional implications, e.g. association with phospholipids or changing the conformation of TM domain(29–31), which might interfere protein-protein interaction, signaling transduction or protein stability(31, 32). In agreement with the latter, our results showed that reducing the level of palm-Cadm4, either by mutation (Cadm4-C347A) or the removal of DHHC3 (DHHC3-KO), possibly promotes dynamin-dependent and Clathrin-mediated Cadm4 internalization from PM and schedules for degradation via autophagy-lysosome path (Fig. 3G and Fig. 5J), which in return regulates the level of Cadm4 on PM. Surprisingly, palmitoylation on other proteins might have opposite effect. It was shown that AMPA receptors are also palmitoylated at the TM/cytosol boundary, upon palmitoylation, AMPA receptors decrease its interaction with cytoskeleton and therefore tend to be internalized and dissociate from PM(25). Additional evidences demonstrated that palmitoylation at such site is involved in diversified regulations e.g. subcellular trafficking or protein-protein interaction etc.(24, 25, 27, 29), suggesting that the roles of palmitoylation at TM/cytosol boundary might vary depending on the proteins investigated but the cysteine residue locates at the TM/cytosolic boundary might be a hotspot for protein palmitoylation.

Interfering Cadm4 palmitoylation, either in Cadm4-KI or DHHC3-KO mice, induces similar but severe myelination abnormalities in CNS, characterized by loss of myelination, myelin infoldings, vacuolation and hypermyelination etc. (Fig. 4B and Fig. 6B), resembling part of the Charcot-Marie-Tooth (CMT) neuropathy(2, 5, 11, 20), suggesting that Cadm4 might be a key substrate of DHHC3 from the perspective of regulating myelination although this cannot rule out other possibilities. It might be logic to understand the loss-of-myelination phenotype, as less-palmitoylated Cadm4 reduces its level on PM (Fig. 4L, Fig. 5E and Fig. 6K), weakens its interactions with Cadm2/3 and Mag (Fig. 4M–4N and Fig. 6L–6M) and hence inhibits myelination or leads to detachment of the axon from the myelin sheath, a scenario was demonstrated constantly in circumstances where either their expression levels or protein-protein interactions were manipulated(5, 7, 8, 11, 20, 33). Apart from that, other myelin abnormalities as thickened myelin sheath in DHHC3-KO/Cadm4-KI mice might be interpreted by altered molecular signaling in guiding myelination. It has been suggested that myelination downstream of erbB2 is controlled by local phosphoinositide production(34, 35), the axonal signaling molecule neuregulin1 type III(36, 37) stabilizes Dlg1 and PTEN complexes by inhibiting their protein degradation, PTEN dephosphorylates PtdIns3,4,5 and antagonizes AKT/PI3K activation (by reducing AKT phosphorylation) downstream of erbB2(38). Notably, the loss of PTEN in myelinating Schwann cells leads to myelin overgrowth as focal hypermyelination and tomaculous neuropathy(35). Furthermore, it was reported that Cadm4 recruits Par3, a multiple-PDZ domain containing protein, to the Schwann cell adaxonal membrane(39), and Par3 interacts with PTEN and regulates cell polarity(40). Interestingly, we did show that AKT phosphorylation is upregulated in the brains of DHHC3-KO and Cadm4-KI as compared to that of the control mice (Fig. S6A–6B), hinting that AKT signaling might be involved in promoting the overgrowth of myelin sheath. However, the most part of the regulatory machinery relating palm-Cadm4, AKT signaling and diversified myelin abnormalities still remains elusive and invites future studies.

Taken together, our findings reveal a regulatory mechanism that the stable localization as well as the level of Cadm4 are regulated by protein palmitoylation, which might provide implications for the neurological diseases as CMT where Cadm4 expression and myelination is abnormally modulated.

## Acknowledgements

We thank Dr. Ying-Chun Hu, Yun-Chao Xie and Peng-Yuan Dong for their professional technical assistance in EM sample preparation and image analysis at the Core Facilities of School of Life Sciences, Peking University. This work was supported by the National Natural Science Foundation of China (Grant No. 32100776 to Y.L.C., No. 31770824 to E.Y.K., No. 81471595 to Y.M.L. and No. 81671388 to S.Q.). The genetic modification of cell lines and mice were supported by 111 program (D20036).

## Author contributions

Y.L.C., J.L.Z., C.C.N., Y.J.L., F.J.R., X.Z.C. and J.J.L. performed experiments and analyzed data; C.Y.Y. provided technical support and data analysis; J.L.Z., T.H.L. and Y.M.L. constructed the Cadm4-KI and DHHC3-KO mice models and joined the discussion; C.H.W., S.Q.Q. and X.H.K. provided technical advice about experimental design and joined the discussion; Y.L.C., J.L.Z. and E.Y.K. initiated the study, designed experiments and wrote the manuscript.

## Data Availability Statement

All data generated in this study are included in the manuscript and supporting files: Figures-source data 1-9 and the MDAR Checklist.

## Material Availability Statement

Materials used in this study are available from the corresponding author on reasonable request.

## Materials and Methods

| Materials | SOURCE | IDENTIFIER |
| --- | --- | --- |
| Antibodies |  |  |
| Anti-Cadm4, rabbit polyclonal | abcam | Cat#ab69605<br>RRID: AB-2259398 |
| Anti-Cadm2, rabbit polyclonal | abcam | Cat#ab64873<br>RRID: AB-1140852 |
| Anti-NECL-1, rabbit polyclonal | abcam | Cat#ab69604<br>RRID: AB-1267900 |
| Anti-Sodium Potassium ATPase, rabbit monoclonal | abcam | Cat#ab76020<br>RRID: AB-1310695 |
| Anti-HSP90, rabbit polyclonal | Cell Signaling | Cat#4874S<br>RRID: AB_2121214 |
| Anti-LAMP1, rabbit polyclonal | Protein-tech | Cat#21997-1-AP<br>RRID: AB_2878966 |
| Anti-Myelin Basic Protein, mouse monoclonal | abcam | Cat#ab62631 RRID: AB-956157 |
| Anti-MAG, rabbit monoclonal | Cell Signaling | Cat#9043S RRID: AB-2665480 |
| Anti-Clathrin heavy chain, rabbit polyclonal | abcam | Cat#ab21679 RRID: AB-2083165 |
| Anti-MEK1 (phosphor S298), rabbit monoclonal | abcam | Cat#ab96379<br>RRID: AB-10678572 |
| Anti-MEK1, rabbit monoclonal | abcam | Cat#ab32091<br>RRID: AB_776273 |
| Anti-AKT, rabbit monoclonal | Cell Signaling | Cat#4691S<br>RRID: AB_915783 |
| Anti-p-AKT (S473), rabbit monoclonal | Cell Signaling | Cat#4058S<br>RRID: AB_331168 |
| Anti-MAPK, rabbit monoclonal | Cell Signaling | Cat#4695S<br>RRID: AB_390779 |
| Anti-p-MAPK (Thr202/Tyr199), rabbit polyclonal | Cell Signaling | Cat#9101S<br>RRID: AB_331646 |
| Anti-GAPDH, mouse monoclonal | Abclonal | Cat#AC033<br>RRID: AB_2769570 |
| Anti-Actin, mouse monoclonal | abcam | Cat#ab7817<br>RRID: AB_262054 |
| Anti-IgG (H+L) Alexa Fluor Plus 647 conjugated, rabbit | Invitrogen | Cat#A21245<br>RRID: AB_142134 |
| Anti-IgG (H+L) Alexa Fluor Plus 594 F(ab')2-conjugated, rabbit | Invitrogen | Cat#A11072<br>RRID:AB_2534116 |
| Anti- IgG (H+L), HRP Conjugate, mouse | Protein Biotechnologies | Cat#PMS301 |
| Anti- IgG (H+L), HRP Conjugate, rabbit | Protein Biotechnologies | Cat#PMS302 |
| Chemicals, peptides, and recombinant proteins |  |  |
| Cycloheximide (CHX) | MedChemExpress | Cat#HY-12320 |
| Chloroquine (CHQ) | Sigma | Cat# |
| 3-Methyladenine | MedChemExpress | Cat#HY-19312 |
| Mg132 | MedChemExpress | Cat#HY-13259 |
| Dynasore | MedChemExpress | Cat# HY-15304 |
| 2-bromopalmitate (2-BP) | Sigma-Aldrich | Cat#238422 |
| Paraformaldehyde | Sigma-Aldrich | Cat#158127 |
| Glutaraldehyde solution | Sigma-Aldrich | Cat#G7651 |
| N-Ethylmaleimide NEM | Sigma-Aldrich | Cat#04259 |
| Hydroxylamine hydrochloride (HA) | Sigma-Aldrich | Cat# 431362 |
| Sepharose 6B | GE | Cat# 17-0420-01 |
| RIPA | Beyotime Biotechnology | Cat# 04693132001 |
| Proteinase inhibitor cocktail | Roche | Cat# 04693132001 |
| mPEG | Nanocs | Cat#PG1-ML-5K |
| Anti-DYKDDDDK Affinity resin | Sigma-Aldrich | Cat# A2220 |
| Cathepsin B (hoechst) | ImmunoChemistry Technologies | Cat# 937 & 938 |
| Glutamax | Gibco | Cat#35050-061 |
| Sodium Pyruvate | Gibco | Cat#11360-070 |
| Penicillin Streptomycin | Hyclone | Cat#SV30010 |
| MEM/EBSS | Hyclone | Cat#SH30024.01 |
| 0.05% Trypsin-EDTA | Gibco | Cat#25300-062 |
| Lipo3000 | Invitrogen | Cat#L3000-008 |
| Opti-MEM | Invitrogen | Cat#31985-070 |
| Poly-D-lysine Hydrobromide | Sigma-Aldrich | Cat#P0671 |
| FBS | Gibco | Cat#10099-141 |
| EMBed 812 | SPI | Cat# 90529-77-4 |
| Critical Commercial Assays |  |  |
| RNeasy Plus Mini Kit | QIAGEN | Cat# 74134 |
| PrimeSCRIPT RT Master Mix | TAKARA | Cat# RR036A |
| TB Green Premix Ex Taq II | TAKARA | Cat# RR820A |
| QIAprep Spin Miniprep Kit | QIAGEN | Cat# 27106 |
| Experimental models: cell lines |  |  |
| 293T | ATCC | Cat#CRL-1573 |
| N2a | ATCC | CCL-131 |
| N2a-zDHHHC3-KO | This paper | N/A |
| Experimental models: organisms/strains |  |  |
| Mouse: C57BL/6 | Beijing Vital River Laboratory Animal Technology Co., Ltd | N/A |
| Mouse: Cadm4-C347A-KI mice | This paper | N/A |
| Mouse: zDHHHC3-KO mice | This paper | N/A |
| Recombinant DNA |  |  |
| pCMV-Cadm4-flag | This paper | N/A |
| pCMV-C339A-flag | This paper | N/A |
| pCMV-C347A-flag | This paper | N/A |
| pCMV-C339/347A-flag | This paper | N/A |
| pCMV-Cadm4-GFP | This paper | N/A |
| pCMV-C347A-GFP | This paper | N/A |
| DHHCS | A gift from Dr. Fukata | N/A |
| pCMV-RFP-LAMP1 | addgene | Plasmid 1817 |
| pCMV-RFP-Rab5 | addgene | Plasmid 14437 |
| pCMV-RFP-Rab7 | addgene | Plasmid 14436 |
| Software and algorithms |  |  |
| ImageJ | NIH | Version 1.50 |
| Photoshop CS6 | Adobe Systems | Version 13.0 x64 |
| Prism | Graph Pad | Version 7 |
| Others |  |  |
| Leica TCS SP8 STED | Leica | N/A |
| HT7700 | HITACHI | N/A |
| Leica UC7 | Leica | N/A |
| Ultra 35° | Daitome | N/A |
| Olympus CKX53 | Olympus | N/A |

### Mouse Models

All animals are kept in SPF environment which is a temperature-controlled room with a 12h light/dark cycle, water and food are adequately supplied. All animal procedures were performed according to guidelines approved by the committee on animal care at Xinxiang Medical University. For all experiments, both male and female littermates were randomly assigned and the principal of double-blind was adopted in all experiments and the following data analysis.

### Generation of Cadm4-KI mice

C57BL/6 (B6) mice were purchased from Beijing Vital River Laboratory Animal Technology. Fertilized B6 eggs were collected and injected via microinjection system as previously described with little modification. In brief, Cas9 mRNA and sgRNA (TCACTTACCCTTCTGTCGGACGG, TGGTGCTCCGTCCGACAGAAGGG) were produced by using in-vitro transcript (IVT) kits, the single-stranded oligonucleotide donor DNA (ssODN) was synthesized by Bioligo company (Shanghai, China), all these components were mixed and microinjected into the cytoplasm of fertilized eggs. Injected eggs were cultured to two-cell stage and then transferred into ICR foster mice. 20-days later, F0 mice were born and genomic DNA was isolated. The primers: F: TAGAGGCGTGTAGATACCAGGAGTGAA and R: GCTAAGACAGAGTCTATCAATGGAGTCCC were used for genotyping. All animal procedures were performed according to guidelines approved by the committee on animal care at Xinxiang Medical University.

### Generation of DHHC3-KO mice

Similarly, Cas9 mRNA and sgRNA (AACTCTGCCTCCACGAGGCA GGG, CCGACACAGTTGTTGACCCA AGG) were produced by using in-vitro transcript (IVT) kits, all these components were mixed and microinjected into the cytoplasm of fertilized eggs. Injected eggs were cultured to two-cell stage and then transferred into ICR foster mice. 20-days later, F0 mice were born and genomic DNA was isolated. The primers: F: TGATGGTTACCCCTGCTGAA and R: TTCTCAACAGCAACCAAGACT were used for genotyping.

### Generation of DHHC3-KO cells in N2a

As N2a is a mouse cell line, we applied the same knockout strategy for N2a as carried out in DHHC3-KO mice. Briefly, two guide RNAs (AACTCTGCCTCCACGAGGCAGGG, CCGACACAGTTGTTGACCCAAGG) were synthesized and cloned into pX458 vector with EGFP which enabled single cell sorting. N2a cells were co-transfected with plasmids of px458-sgRNA1 and px458-sgRNA2 by Lipofectamine 3000 reagent with the reduced serum medium Opti-MEM. Two days later, single cell was sorted by BD FACSAria™ Fusion and then cultured until the formation of colonies. Individual colonies were screened by PCR and sanger sequencing was performed for further verification. N2a was maintained in MEM medium (Gibco, USA) supplemented with 10% fetal bovine serum (Gibco, USA), 100 μg/ml streptomycin, 100 U/ml penicillin, 1mM sodium pyruvate and 1% glutamine (100×, Gibco), and incubated in 5% CO_2_ at 37°C.

### Plasmids

The plasmids expressing mouse-origin Cadm4-Flag, Cadm4-GFP, APT1-Flag, APT2-Flag, PPT1-Flag, PPT2-Flag and ABHD17a-Flag were constructed. The cDNA for the mouse Cadm4, APT1, APT2, PPT1, PPT2 or ABHD17a was obtained by RT-PCR which was then subcloned into pUC19 as cDNA donors. Specific primers were synthesized for subcloning individual cDNA into pCMV3-C-Flag for mammalian cell expression by In-Fusion cloning method. The Cadm4-C339A-Flag, Cadm4-C347A-GFP, Cadm4-C347A-Flag, Cadm4-C339/347A-GFP and Cadm4-C339/347A-flag mutants were generated by site-directed mutagenesis PCR reaction. Other plasmids as RFP-Rab5 (Plasmid 14437), RFP-Rab7 (Plasmid 14436) and RFP-Lamp1 (Plasmid 1817) were obtained from Addgene. All plasmids of Ha-DHHCs were a gift from Dr. Fukata. All constructs were verified by sequence analysis.

### Cell culture, transfection and treatments

HEK-293T (CRL-11268) and N2a (CCL-131) were obtained from ATCC. HEK-293T and N2a were grown in DMEM high glucose medium (Gibco, USA) and MEM (Hyclone) respectively, supplemented with 10% fetal bovine serum (Gibco, USA) containing 100 μg/ml streptomycin and 100 U/ml penicillin at 37 °C in a 5% CO_2_ incubator. All cell lines were transfected using Lipofectamine 3000 reagent for 24 hr (Invitrogen) with the reduced serum medium Opti-MEM (Life Technologies). The follow reagents were used for cells treatments: Dimethyl sulfoxide (DMSO) (Sigma-Aldrich, Cat# D8418), 50μM 2-Bromohexadecanoic acid (2-BP) (Sigma-Aldrich, Cat# 238422), 100μM Cycloheximide (CHX, MedChemExpress, HY-12320), 25μM Mg132 (MedChemExpress, HY-13259), 50μM Chloroquine (CHQ, Sigma, C6628), 3mM 3-Methyladenine(3-MA, MedChemExpress, HY-19312) and 80μM dynasore (MedChemExpress,HY-15304). All drug experiments were started at 12hr after cells were transfected. Cells were exposed to fresh MEM complete medium containing drugs or control DMSO for the indicated time and then washed with PBS for 3 times.

### Acetyl-Resin-assisted capture assay (Acyl-RAC)

RAC assay was performed as previously described with minor modification (NEM was used to replace MMTS). Briefly, mice brain/cells were lysed in lysis buffer (20 mM Tris pH=7.5, 150 mM NaCl, 1% Triton X-100) containing protease inhibitor cocktail (Roche). Lysate was sonicated and incubated at 4 °C for 30 min while rotating, which was then cleared by centrifugation at 15,000 g for 10 min at 4 °C, the supernatant was havested. The amount of protein was quantified with a bicinchononic acid (BCA) assay Kit (Cat#P0009, Beyotime). Protein lysate (1 mg) was diluted to 1 mg/ml with blocking buffer (100 mM HEPES, 1 mM EDTA, 2.5% SDS, 50 mM NEM, pH 7.5) and kept at 50°C for 60 min with agitation. NEM was removed by four sequential 70% acetone precipitations, pellet was resuspended in 600 μl binding buffer (100 mM HEPES, 1 mM EDTA, 1% SDS, pH 7.5). Samples were divided into two equal parts and added with 50 μl pre-equilibrated thiopropyl Sepharose 6B (Cat#17-0420-01, GE Healthcare) respectively, one part was treated with 40 μl 2 M hydroxylamine (HA+) pH 7.0 (to cleave thioester linkage) and the other part was incubated with 40 μl 2 M NaCl (negative control, HA-), 20 μl of each supernatant was taken as ‘input’. Cleavage and capture were carried out on a rotator at room temperature for 4 hr. Resins were washed five times with binding buffer, eluted with 40 μl Leammli loading buffer (2.1% SDS, 66 mM Tris-HCl (PH7.5), 26% (W/V) glycerol, 50 mM DTT) on a shaker at 42°C for 15 min and subjected for Western blot analysis.

### mPEG-labeling assay

mPEG-labeling was performed as described previously. Cultured cells/tissues were lysed with 4% SDS in TEA buffer (50 mM Triethanolamine, 150 mM NaCl, pH=7.3) supplemented with protease inhibitor cocktail (Roche). Total 500μg protein was treated with 10mM TCEP for 30min and then incubated with 25mM NEM for 2 h at room temperature to block unmodified cysteine residues. NEM blocking was terminated by 3-times methanol-chloroform-H2O precipitation (4/1.5/3) and dried in a speed-vacuum (Centrivap Concentrator, Labconco) after the last precipitation to remove NEM completely. The pellet was resuspended in 100 μl TEA buffer containing 4% SDS and incubated at 37 degree for 10 mins with gentle agitation. Next, the samples were treated with 1M neutralized NH2OH at room temperature for 1h to cleave the thioester linkage formed between cysteine residue and palmitate, which were then subjected for methanol-chloroform-H2O precipitation and resuspended in 30μl TEA buffer containing 4% SDS and 4mM EDTA at 37 degree for dissolving. For mPEG-maleimide alkylation, additional solution containing 0.2% Triton X-100, 90μl TEA buffer and 1.33 mM mPEG-Mal (final concentration at 10 mM) was added and incubated at room temperature for 2h. Last, the reaction was stopped by a final methanol-chloroform-H2O precipitation and resuspended in laemmli buffer (Bio-Rad) for WB analysis.

### Immunofluorescence staining and imaging

N2a cells were plated onto poly-D-lysine-coated coverslips and transfected as described with indicated plasmids. At 24 hours post-transfection, the cells were fixed with 4% (W/V) paraformaldehyde (Electron Microscopy Sciences, Cat#15710). After blocking in 3% BSA in PBS, cells were stained (without permeabilization) with primary antibody Cadm4 (abcam, Cat# ab69605, 1:500), Na/K ATPase (Abcam, Cat# ab76020, 1:300), Flag-M2 (sigma, F3165, 1:1000) and then incubated with secondary antibodies Anti-Rabbit IgG (H+L) Alexa Fluor Plus 488 conjugated (Invitrogen, Cat#A11070, 1:1000), Anti-Rabbit IgG (H+L) Alexa Fluor Plus 594 conjugated (Invitrogen, Cat#A11072, 1:1000) and Alexa Fluor Plus 647 conjugated (Invitrogen, Cat#A21245, 1:1000), washed, and mounted onto slides with Fluoromount-G (Electron Microscopy Sciences, Cat#17984-25). The fluorescence images were taken with a Stimulated Emission Depletion microscopy (Leica TCS SP8 STED).

### Plasma membrane fractionation

Plasma membrane fractionation was performed as described previously (Li etal, 2017), with minor modifications. N2a cells/tissues were homogenized in HME buffer (10mM HEPES, 1mM MgCl2, 1mM EDTA, PH7.4) supplemented with protease inhibitor cocktail (Roche). The homogenates were then subjected for Freeze/Thaw cycle (freeze in liquid nitrogen, thaw at 37°C) for 5 times, and then sonicated 10 seconds on/off for 1 min on ice using a microsonicator (Scientz, UP-250). The homogenates were centrifuged at 900g for 10 min to remove cell debris and nuclei, the supernatant was collected and centrifuged at 20000g for 40min at 4 degree and the supernatant was collected as cytosol fractions. The pellets (P1) were resuspended in HEM buffer (with protease inhibitor) and centrifuged at 10000g for 30min, 4 degree. The pellets (P2) were collected as subcellular organelle membrane fractions, and the supernatant was subjected for last centrifugation at 20000g for 40min, 4 degree, the pellets (P3) were collected as plasma membrane fractions (majority). For verification, all fractions were examined by Western blot and their corresponding marker proteins (HSP-90 and Na/K ATPase were used as markers for cytosol and plasma membrane fractions respectively).

### Western blot analysis and antibodies

Samples were separated in standard SDS-PAGE gels and transferred to Immobilon-P PVDF membrane (Pore size 0.2 μM, EMD Millipore). The membrane was then blocked in 5% (w/v) skimmed milk in TBS containing 0.1% (v/v) Tween-20 (TBST) for 90 min. After blocking the membranes were washed in TBST and incubated with primary antibody for overnight at 4°C. After washing with TBST, the membranes were incubated with a suitable horseradish peroxidase (HRP) labeled secondary antibody and signals were detected with an ECL kit (Tanon). The following primary antibodies were used: The following primary antibodies were used: Cadm4 (Abcam, Cat# ab69605, 1:1000), Cadm2 (Abcam, Cat# ab64873, 1:1000), Cadm3 (Abcam, Cat# ab69604, 1:1000), Na/K-ATPase (Abcam, Cat# ab76020, 1:20000), HSP90 (Cell Signaling, Cat# 4874S, 1:1000), MAG (Cell Signaling, Cat# 9043S, 1:1000), Flag (Abclonal, Cat# AE024, 1:5000), Actin (Abcam,Cat# ab7817 1:10000) and Gapdh (Abclone, Cat# AC033, 1:5000). The following secondary antibodies were used for immunoblot: Goat Anti-Rabbit IgG (H+L), HRP Conjugate (Protein Biotechnologies, Cat# PMS302, 1:5000) and Goat Anti-Mouse IgG (H+L), HRP Conjugate (Protein Biotechnologies, Cat# PMS301, 1:5000).

### Immunoprecipitation

Briefly, cell/tissue lysates were incubated with corresponding antibody (anti-Cadm4, 1:200) or IgG and kept for rotation at 4 degree overnight. The next day, protein A/G beads, which was pre-equilibrated in PBS (pH 7.4), was added and kept for additional 2h at 4 degree in rotating. The beads were then collected at 1000 g for 2 min and washed at least 3 times with lysis buffer coupled with protease inhibitor cocktail (Roche). After last wash and centrifugation, the beads were eluted with 1 × SDS-PAGE Loading Buffer (Beyotime, Cat#P0015L), heated at 100 degree for 5 mins and subjected for Western blot analysis.

### Sample preparation and imaging for Electron Microscopy

Anesthetized mice were perfused with normal saline followed by perfusion with freshly prepared fixative containing 4% paraformaldehyde and 2.5% glutaraldehyde supplied with 0.1 M sucrose in 0.1 M phosphate buffer (PB), pH 7.4. The brains were chopped to 1 mm^3^ blocks, kept for 2 h at room temperature, and then stored overnight at 4°C. After washing four times with 0.1 M PB, the samples were post-fixed in 2% OsO4 /1.5% potassium ferrocyanide for 1 h at room temperature. Then the samples were rinsed several times in ddH2O and en bloc stained with 2% aqueous uranyl acetate overnight at 4 °C. Following several washes in ddH2O, specimens were dehydrated in a series of ethanol solution (30%, 50%, 70%, 85%, 95%, 100%, 100%; 10 min each) and replaced with pure acetone two times for 10 min each. Tissue samples were gradually permeated and embedded with EMbed 812 resin (Electron Microscopy Sciences). Resin blocks were sectioned with a diamond knife (ultra 35°, Diatome, Switzerland) using an ultramicrotome (UC7, Leica Microsystem) and collected in copper grids with a single slot. Ultra-sections (70 nm) were stained with uranyl acetate and lead citrate and examined under electron microscopes Tecnai G2 Spirit (Thermo Fisher Scientific) or Hitachi HT7700 equipped with a CCD camera (Orius 832, Gatan) at 120kV. EM micrographs were analyzed by Image J (version 1.50) for myelin thickness and axon diameter. G-ratio was calculated by dividing the inner axonal diameter by the outer axonal diameter. Around 300 axons from 3 mice (at around P90) were randomly chose for quantification for each genotype.

### Statistical Analysis

Basic descriptive data are presented as means standard errors of means. For statistical analyses of differences between two groups, paired or unpaired two-tailed Student’s t tests were used where appropriate. For experiments involving more than two groups, one-way ANOVA analysis was carried out. Post-hoc pairwise comparisons, with Bonferroni correction for multiple comparisons, were conducted where appropriate. An α-level of 0.05 was adopted in all instances. All analyses were carried out using SPSS 19 professional software (IBM, USA). Graphs were created using GraphPad Prism (Windows, version 7). None of the samples were excluded from the statistical analysis. Sample sizes referred to the general application of the field and were not statistically predicted.

## Supplemental Information

**Fig. S1.**
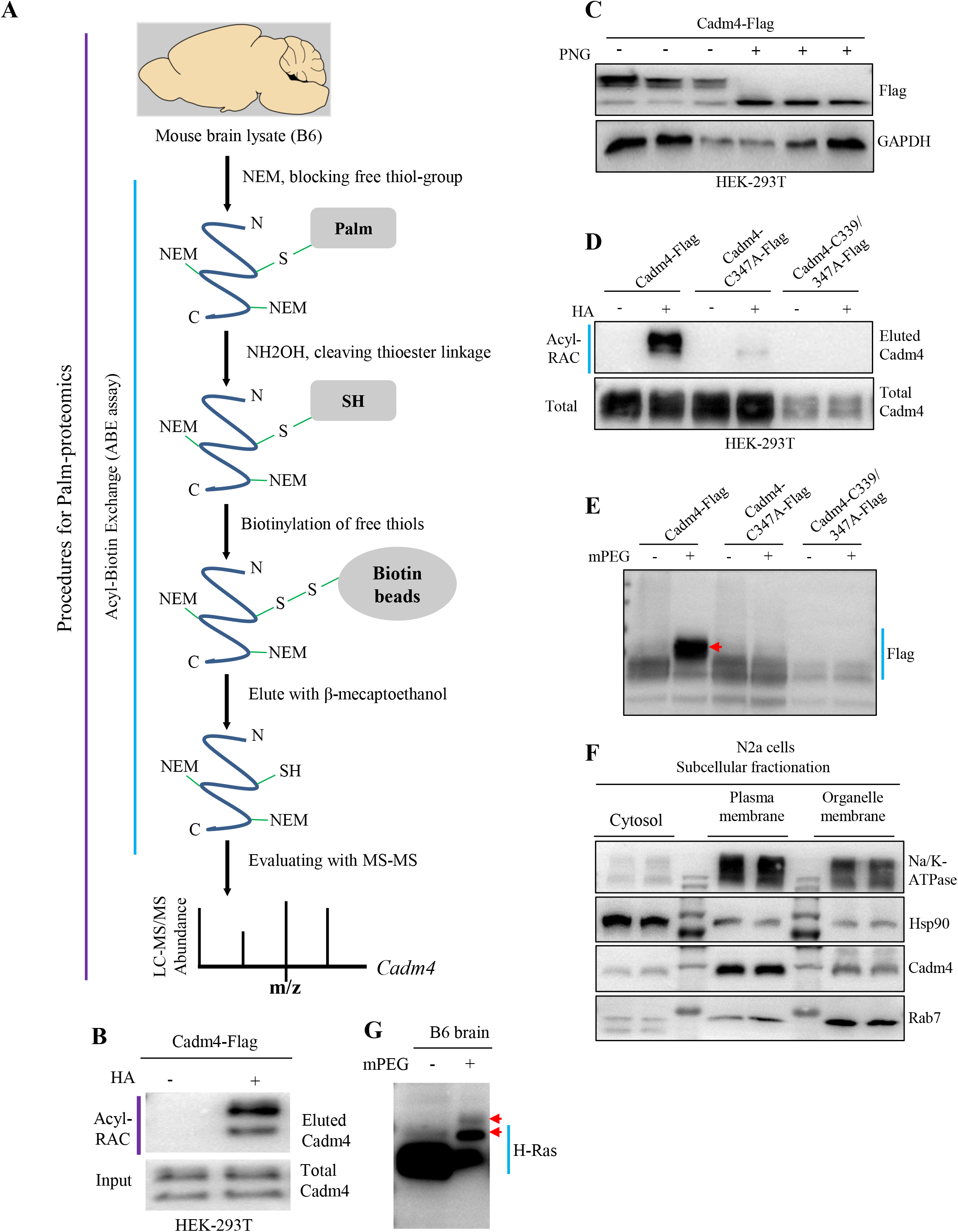
Identification of Cadm4 as palmitoylated protein. **A**, A diagram to show the procedures of palm-proteomics, where Cadm4 was initially identified. **B**, Cadm4 expressed in HEK-293T cells was analyzed for palmitoylation by Acyl-RAC assay. **C**, HEK-293T cells expressing Cadm4 was incubated with or without PNG (an inhibitor of protein glycosylation) for WB analysis. **D**, Cadm4 or its mutants (C347A and C339A/C347A) were expressed in HEK cells and subjected for Acyl-RAC assay. **E**, HEK cells expressing Cadm4 or its mutants (C347A and C339A/C347A) were processed for mPEG-labeling assay. The migrated band is pointed by the red arrow. **F**, Subcellular fractions were prepared from WT N2a cells, HSP-90 is a marker for cytosol proteins, Na/K ATPase is a marker for plasma membrane proteins. **G**, WT brain lysate was processed with mPEG-labeling assay for detecting H-Ras palmitoylation. The mPEG labeling causes the band shift, pointed with red arrows.

**Fig. S2.**
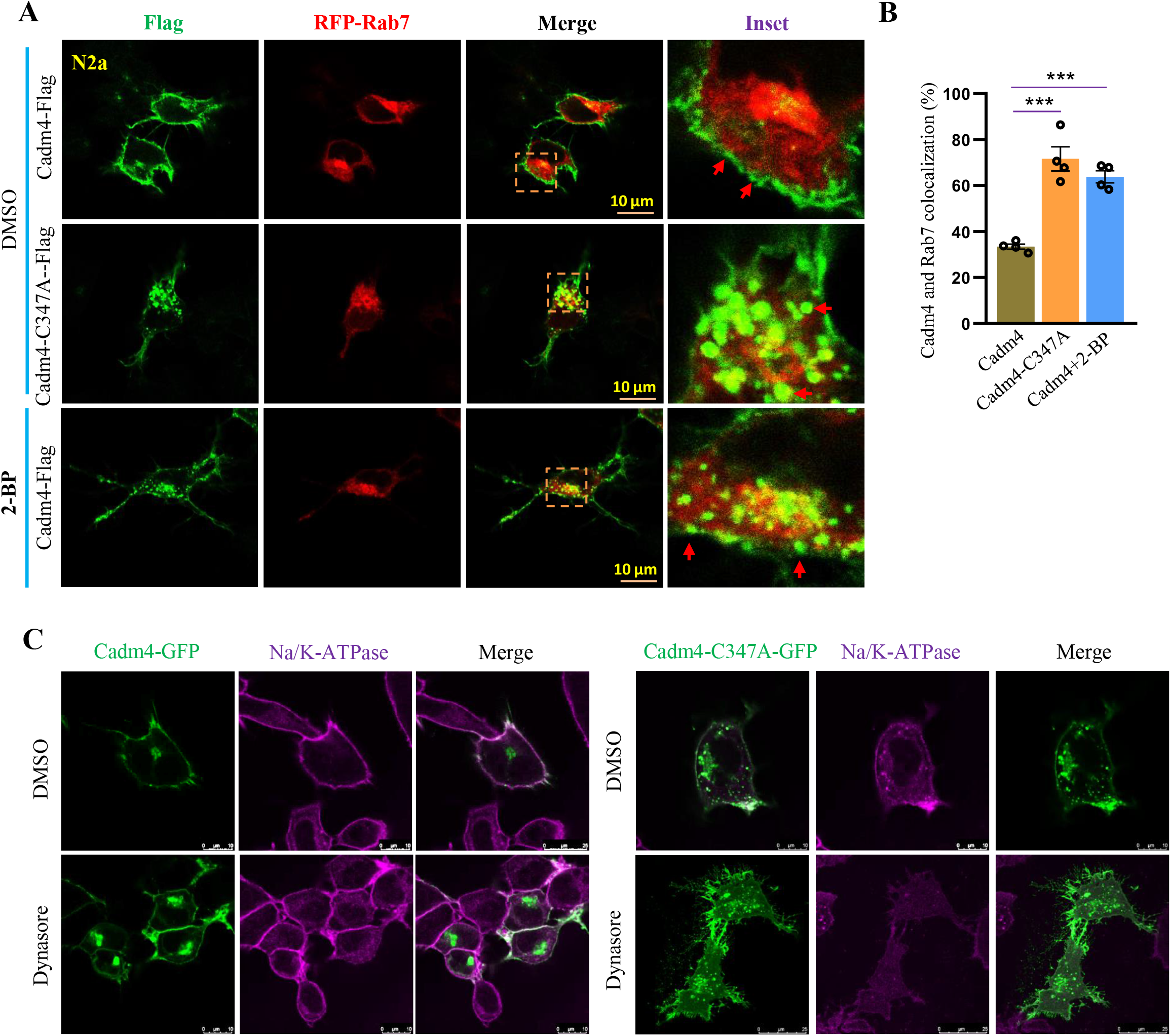
Dynasore is capable of inhibiting Cadm4 internalization. **A**, Cadm4-GFP or Cadm4-C347A-GFP was expressed in N2a cells, treated with DMSO or 2-BP (50 μM) and fixed for immunofluorescence analysis, RFP-Rab7 was used to label the late endosome. **B**, The colocalization rate of Cadm4 and Rab7 was quantified. \*\*\**P*< 0.001, n=4 cells, one-way ANOVA; Bonferroni post hoc test comparing Cadm4 and Cadm4-C347A, \*\*\**P*< 0.001; Cadm4 and Cadm4+2-BP, \*\*\**P*< 0.001. **C**, N2a cells expressing Cadm4 or Cadm4-C347A were incubated with DMSO or Dynasore (80 μM), and fixed for immunofluorescence analysis. Data are mean ± s.e.m.

**Fig. S3.**
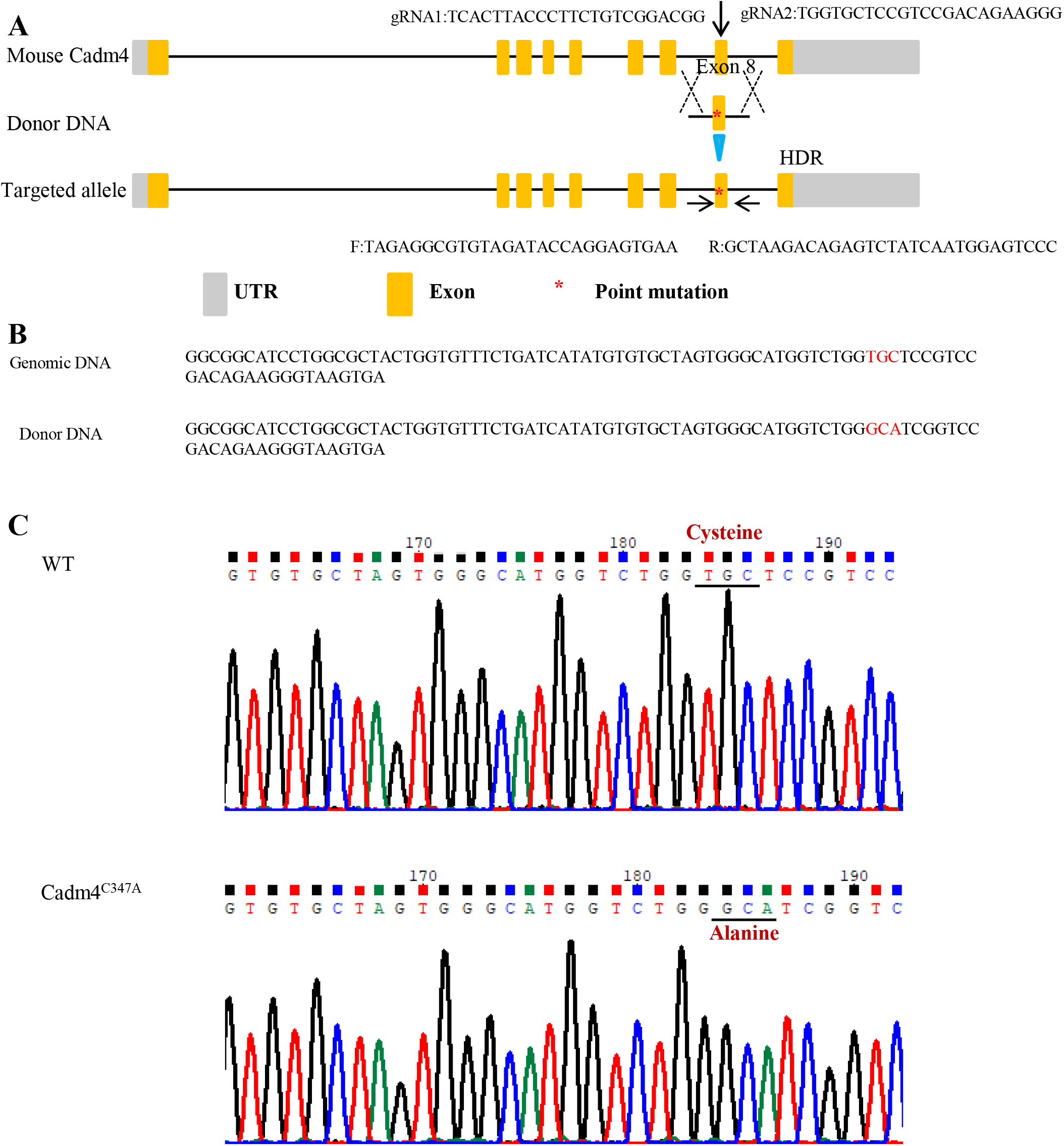
Generation of Cadm4-C347A point mutation mice. **A**, Targeting scheme of the point mutation (C347A) in the 8th exon of mouse Cadm4, the point mutation was introduced into mouse fertilized eggs using Crispr/Cas9 with the HDR donor DNA. Yellow box indicates exons, gray box indicates UTRs. Red star indicates a TGC→GCA mutation in the 8th exon of mouse Cadm4; **B**, The sequences of WT genomic DNA and donor DNA. TGC to GCA (red) was designed to change the amino acid Cys to Ala; **C**, Representative sequencing result of Cadm4-C347A point mutation mice. The mutation TGC to GCA was underlined and the changed amino acid Cys to Ala was in dark-red.

**Fig. S4.**
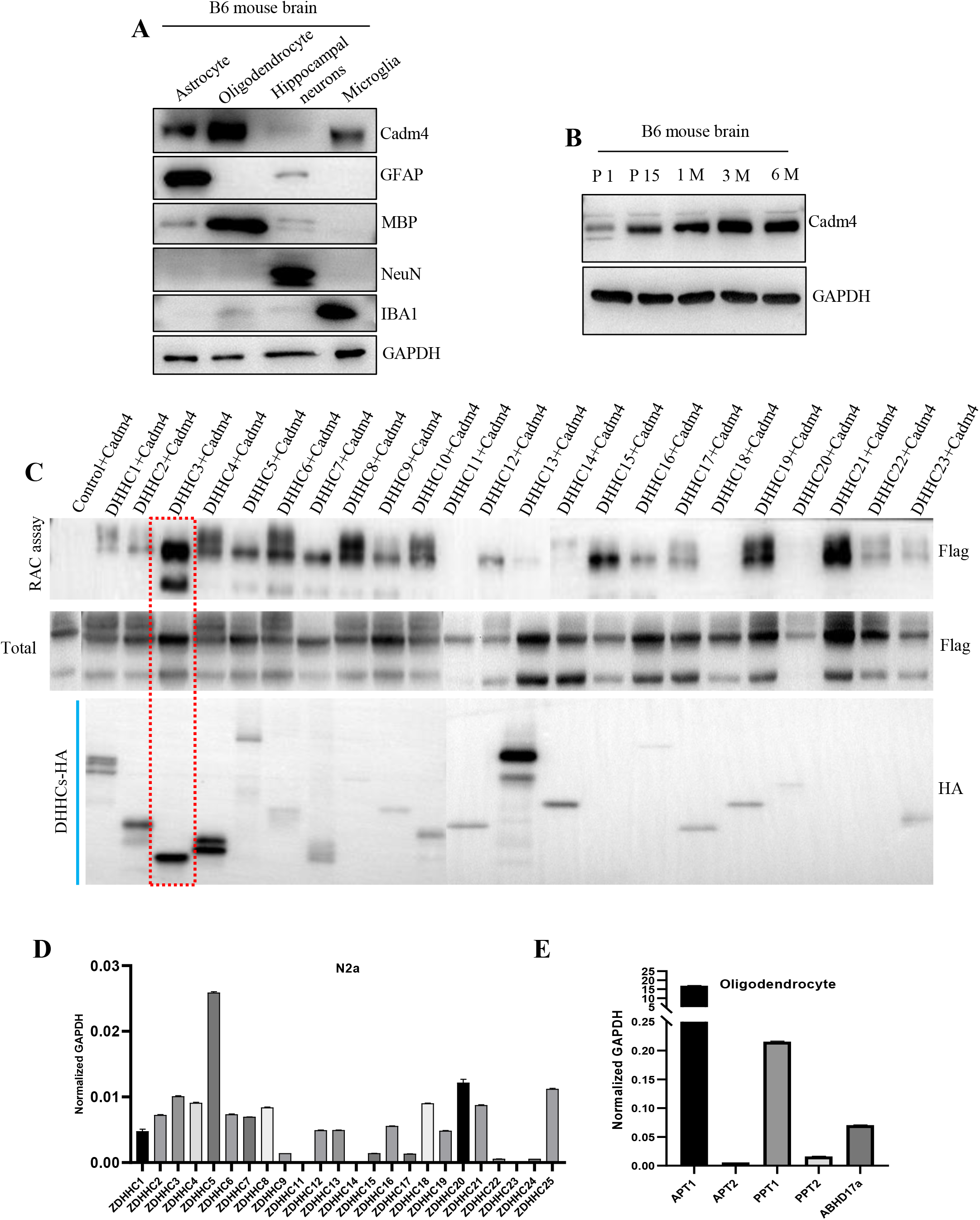
DHHC3 catalyzes the palmitoylation of Cadm4, related to Fig. 6. **A**, Various cell types were isolated and cultured from P0 WT mice, and the protein level of Cadm4 was measured by WB. **B**, Wildtype brain lysates of different developmental stages were collected for measuring the protein level of Cadm4. **C**, All DHHCs were coexpressed with Cadm4 in N2a cells for evaluating the level of palm-Cadm4 by Acyl-RAC. **D-E**, The relative mRNA levels of all DHHCs were quantified by Real-Time PCR in either N2a cells (**D**) or cultured oligodendrocytes (**E**) isolated from P0 WT mice.

**Fig. S5.**
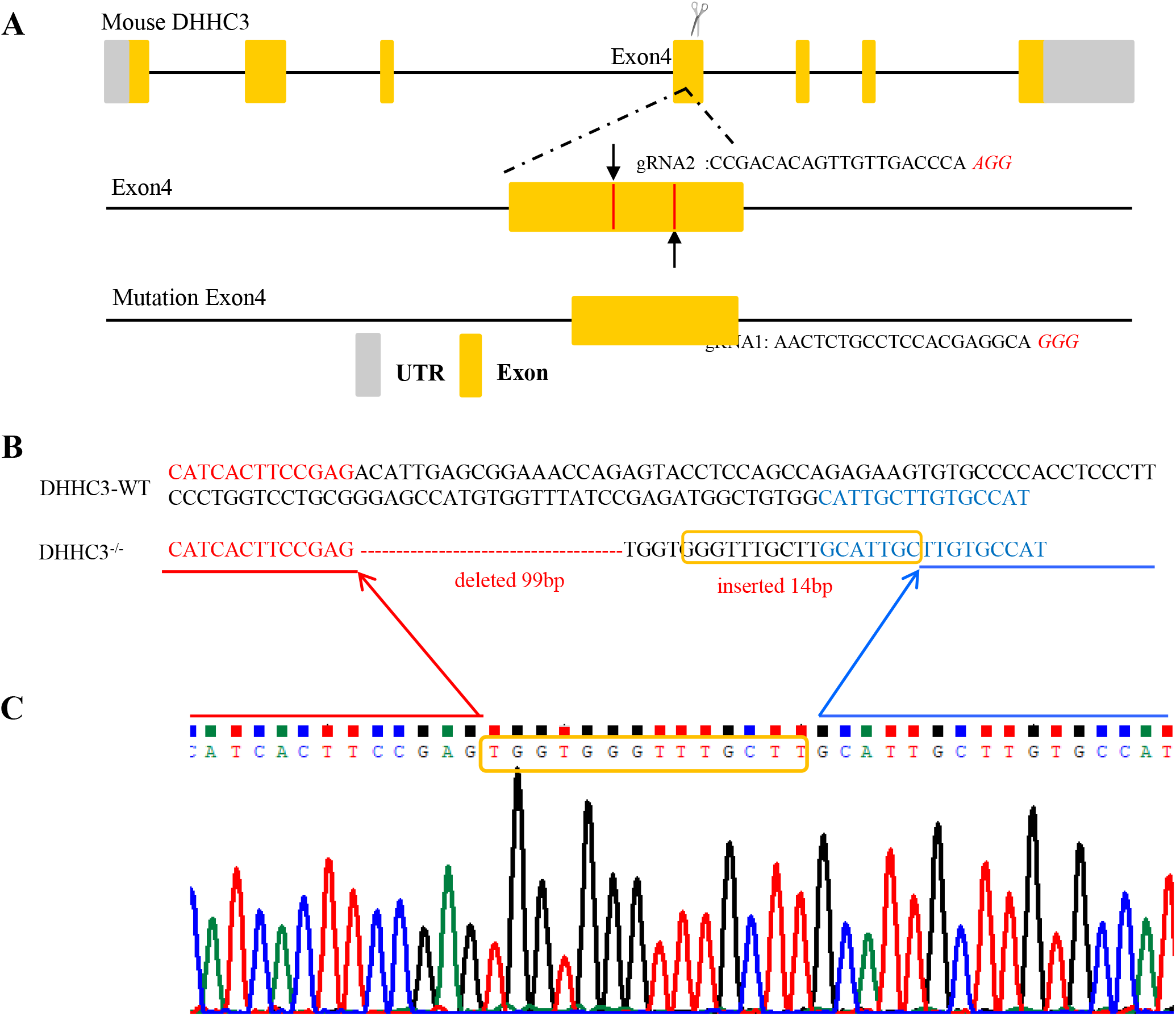
Generation of DHHC3-KO mice. **A**, Targeting scheme of the truncation in the 4th exon of mouse DHHC3, where the enzyme activity center locates. The deletion mutation was introduced into mouse fertilized eggs (C57/B6 background) using Crispr/Cas9 with two gRNA. Red box indicates exons, gray box indicates UTRs. Red lines indicate two cut sites in the 4th exon of mouse DHHC3. **B**, The sequences of WT genomic DNA and mutation DNA. 99bp were deleted from the mutation DNA, but inserted 14 bp, together it caused frameshift and an earlier stopcodon. **C**, Representative sequencing result of DHHC3-KO mice. The insertion was outlined. The evidence at the protein level was not shown due to the fact that the commercially available DHHC3-antibody lacks specificity.

**Fig. S6.**
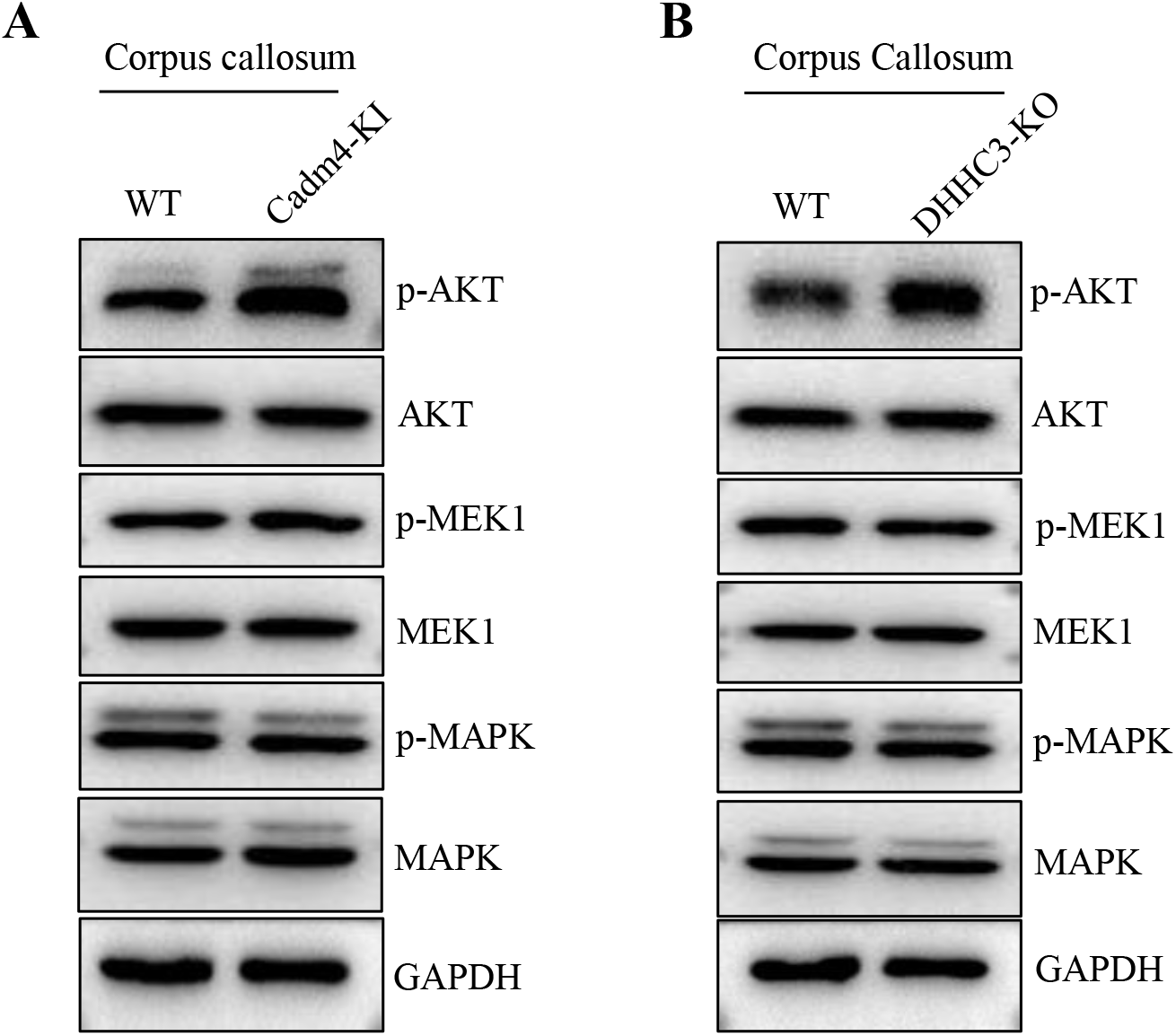
Phospho-AKT is upregulated in the corpus callosum of Cadm4-KI and DHHC3-KO mice. **A-B**, Lysates (prepared with phosphatase inhibitors) from either the corpus callosum of Cadm4-KI (**A**) and DHHC3-KO (**B**) mice were analyzed for the levels of various proteins related to the classical signaling cascades downstream of PI3K.

## Source Data

**Figure 1-source data 1**, uncropped blots for Fig. 1.

**Figure 2-source data2**, uncropped blots for Fig. 2.

**Figure 3-source data3**, uncropped blots for Fig. 3.

**Figure 4-source data4**, uncropped blots for Fig. 4.

**Figure 5-source data5**, uncropped blots for Fig. 5.

**Figure 6-source data6**, uncropped blots for Fig. 6.

**Figure S1-source data7**, uncropped blots for Fig. S1.

**Figure S4-source data8**, uncropped blots for Fig. S4.

**Figure S6-source data9**, uncropped blots for Fig. S6.

